# AlphaFlex: Ensembles of the human proteome representing disordered regions

**DOI:** 10.1101/2025.11.24.690279

**Authors:** Zi Hao Liu, Oufan Zhang, Stefano De Castro, Kunyang Sun, Hamidreza Ghafouri, Omar Abdelghani Attafi, Nicolas L. Fawzi, Silvio C. E. Tosatto, Alexander M. Monzon, Alan M. Moses, Teresa Head-Gordon, Julie D. Forman-Kay

## Abstract

More than two thirds of proteins in the human proteome are predicted to contain intrinsically disordered regions (IDRs), which lack stable folded structure. IDRs are critical for biological regulation and organization, as targets for post-translational modifications, and as mediators of biomolecular condensates. To address the pressing need for better structural models enabling functional insight, we developed AlphaFlex to model fully atomistic conformer ensembles for proteins predicted to have IDRs, modeled in the context of AlphaFold folded domains and an implicit bilayer for transmembrane proteins. The AlphaFlex resource provides conformational ensembles of human proteins from the AlphaFold database with identified IDRs in the Protein Ensemble Database that is mirrored in UniProt. This transformative resource of AlphaFlex ensembles provides physically and biologically relevant full-length models for IDR proteins, including scaffold proteins, those with IDR:folded-domain interactions, regulatory and condensate proteins requiring exposed binding elements, conditionally folding IDRs, and transmembrane proteins containing IDRs.

## INTRODUCTION

Structure has driven functional insight into biology ever since the elucidation of protein α-helical structure^1^ in 1951, followed by years of experimental structure determinations, creating the necessary data for the AlphaFold breakthrough for predicting folded protein structure^2^. Today, the AlphaFold database (AFDB) of predicted protein structures has had a dramatic impact on the biological community, advancing the widely accepted protein structure-function paradigm central to the folded state^2^. However, it is increasingly appreciated that all proteomes also encode intrinsically disordered proteins and regions (IDPs/IDRs, referred to here as IDRs), which do not adopt a well-defined tertiary structure, but instead populate fluctuating and heterogeneous structural ensembles^3–6^. Their highly dynamic nature facilitates diverse biological functions (e.g., transcription, translation, signaling) via a variety of mechanisms including direct protein and nucleic acid binding^7,8^, scaffolding cellular structures and protein complexes^5,9,10^, mediating the formation of biomolecular condensates^11^, exerting pressure through excluded volume^12^, and creating appropriate spacing between folded domains^13,14^. For roughly a third of the residues in the canonical human proteome that are located in IDRs^15^, the AFDB only provides low confidence predictions^2^ that represent IDRs as a single conformational state rather than a conformational ensemble. In general, the single conformations AFDB provides for IDRs often appear as featureless extended curves surrounding clustered folded domains which obscures binding motifs. A further aspect of this is that AlphaFold2 incorrectly predicts confident structures for conditionally folding IDRs (i.e., those that fold upon binding or post-translational modification), estimated to involve about 15% of human IDR residues^16^.

The poor modeling of IDRs and hence also the orientations/proximities of folded domains connected by IDRs presents a major issue, since understanding IDR biology is hampered by the lack of conformational ensembles that effectively represent the heterogenous sampling so critical for their function. Having full-length structural ensembles that correctly represent IDRs in the context of folded domains and transmembrane bilayers is crucial for understanding protein structure-function relationships and providing biologically relevant models. Regulatory IDRs can block catalytic, allosteric or binding sites, with the block “removed” by changes in post-translational modifications or binding to other partners^17^. Conformational ensembles that show the extent of conformational heterogeneity, including multiple potential interactions or lack thereof, provide insights into such regulatory mechanisms. Another example is for conditional folding, where having structural models of the disordered and the folded states is key for understanding how the condition impacts downstream events^16^. An important example is the case of scaffolds and proteins involved in biomolecular condensates. These most often contain multiple folded domains that bind proteins and/or nucleic acids along with IDRs that function as linkers, low-complexity phase-separating regions, and/or modulators of chain solvation ^8,18^. The long “reach” of extended IDRs is essential for facilitating binding to various targets, which may be geometrically constrained, and the conformational heterogeneity is key for exchanging interactions that underlie condensate formation. An example of the need for chain extension required for function is found with the N-terminal IDR (150 aa) of SMC4, a key component of the condensin complex required for condensed chromatin. This SMC4 IDR contains DNA-binding regions and is involved in forming biomolecular condensates that promote proper chromosome compaction, segregation, and genomic stability^7^. The extension of the N-IDR of SMC4 is proposed to enhance the activity of the condensin complex by contacting distant DNA base-pairs. These considerations also apply to the ∼50% of human transmembrane proteins that contain at least one IDR ≥ 30 aa, with these IDRs preferentially located on the cytoplasmic side and enriched for post-translational modifications, facilitating interactions and signaling-related functions^19^.

Hence, beyond AFDB that offers full-length atomistic models focused on folded proteins, a corresponding database of all-atom protein ensembles that represent the IDRs within the context of folded domains is sorely needed. Here, we present the AlphaFlex scalable workflow that leverages AlphaFold’s strengths for predicting the folded domains, while overcoming its known deficiencies by generating all-atom IDR ensembles across the disordered regions. Of the 23,391 proteins from the canonical human proteome in the AFDB^20^, we identify 14,792 proteins (63%) as having an IDR of 15 or more consecutive amino acids using a union of five independent IDR predictions. We use two independent approaches, IDPConformerGenerator^21,22^ (AFX-IDPCG models) and IDPForge^23^ (AFX-IDPForge models), to generate full-length protein ensembles of atomistic conformations including the IDRs and the folded domains, which are made easily accessible in the AlphaFlex database. The AlphaFlex ensembles can be used for hypothesis generation, and as starting structures for molecular dynamics simulations or reweighting based on experimental data, as well as using experimental data during generation using IDPConformerGenerator or IDPForge. Within the set of 14,792 disorder-containing proteins, we identify 2,450 that are annotated as transmembrane proteins and amenable to an AlphaFlex workflow restricting IDR conformational sampling an implicit bilayer. At present there are 9,315 completed protein ensembles, including 231 transmembrane proteins restricted by an implicit bilayer, deposited in the Protein Ensemble Database (PED)^3^ (https://proteinensemble.org) that is mirrored in the UniProt database.

These completed AlphaFlex protein ensembles not only offer a better representation for proteins containing IDRs but also exemplify the full range of IDR complexity, from terminal IDRs through to multiple IDR regions which separate both non-interacting and interacting folded domains, and a range of sequence lengths up to over a thousand amino acids, allowing us to consider their biological implications. Analysis of the AlphaFlex ensembles reveals that IDRs generated within the context of interacting folded domains have statistically different global structural properties compared to previous work that produced IDR ensembles in isolation^21,24^. AlphaFlex ensembles have global structural properties and Cα-Cα distances that are better aligned with the scaffold role of a subset of these proteins than the structures provided by the AFDB^2^. Furthermore, AlphaFlex ensembles contain more fractional α-helical content, especially at longer sequence lengths, than representations of IDRs generated by methods such as CALVADOS^24^, AF-CALVADOS^25^, and, as expected, than AFDB protein regions of low confidence^2^. This helical content can correspond to biological functionality^26,27^, particularly in the context of binding. Additionally, the AlphaFlex structural ensembles allow for accessibility to protein binding sites and regions of post-translational modifications, sites that are more occluded by the AFDB models. The AlphaFlex ensembles provide a more realistic representation compared to AFDB models after benchmarking against full-length SAXS and NMR chemical shift experimental data, while providing meaningful interpretations regarding biological function that we illustrate for proteins whose IDRs regulate cell morphology^28^, transcription^29^ and translation^26^, phase separation^30^, and RNA stability^31^. AlphaFlex ensembles are publicly available as a workflow tool and through deposition in the PED^3^ as a transformative resource for the biological community, paralleling the previous impact of the AFDB.

## RESULTS

### AlphaFlex workflow for generating atomistic ensembles of proteins with IDRs

A diagram representing the AlphaFlex workflow is given in Fig. 1. Stage 1 of the workflow assigns each residue in a protein sequence as part of an IDR or folded domain. The predicted local distance difference test (pLDDT) metric provided by the AFDB has been used as an indicator of intrinsic disorder^16,32^. However, pLDDT is not intended to be a disorder predictor and it is incorrectly confident in predicting structure for cases of conditionally folding IDRs^16^. Therefore, we define an IDR as 15 or more consecutive amino acids predicted to be disordered by a union of 5 metrics, including pLDDT < 70 and disorder prediction from 4 *bona-fide* predictors of disorder: metapredict^33^, flDPnn^34^, ADOPT^35^, and SPOT-Disorder^36^. Statistics for the number of IDRs from each metric can be found in Supplementary Fig. S1. A folded domain region is defined by at least 5 consecutive residues that are not classified as disordered by the union of the 5 metrics. Note that “folded domain region” does not need to encompass a full folded domain, with folded elements potentially interspersed with disordered loop regions.

**Fig. 1.**
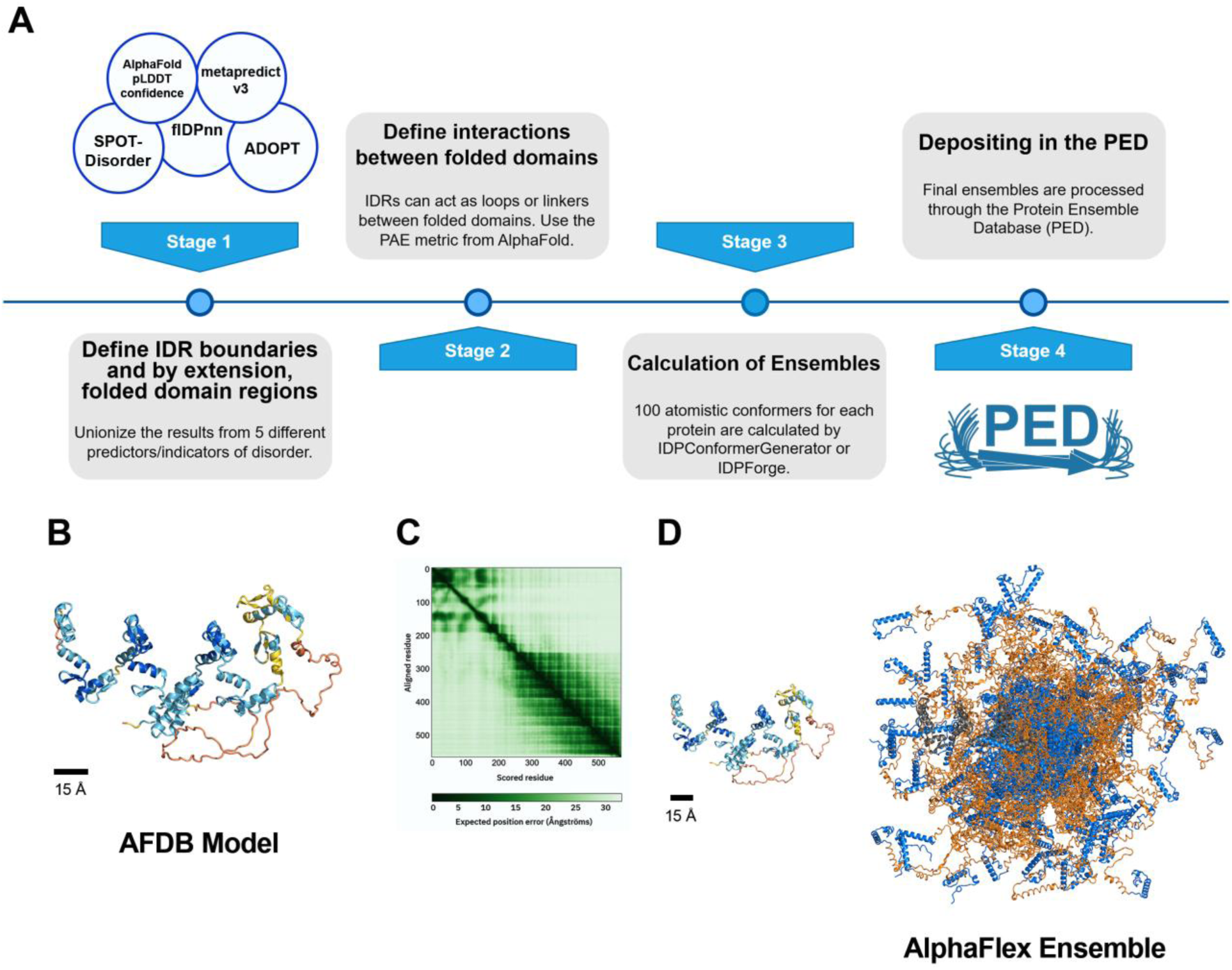
AlphaFlex workflow summary. (**A**) Stages of the AlphaFlex workflow. In Stage 1, IDRs are defined as the union of 5 different predictors/indicators of disorder. Stage 2 defines whether two confidently predicted folded domains or segments are interacting based on AFDB PAE (predicted alignment error). Calculations of atomistic ensembles are done in Stage 3 using information from Stages 1 and 2, with either IDPConformerGenerator^21,22^ or IDPForge^23^. In Stage 4, ensembles are processed and deposited into the Protein Ensemble Database^3^. (**B**) AFDB structural prediction of Zinc finger protein 675 (AF-Q8TD23-F1-v6), colored by pLDDT score values, with yellow and orange representing low (50 < pLDDT < 70) and very low (pLDDT < 50) confidence, respectively, an indicator of disorder (pLDDT < 70), and light blue (70 < pLDDT < 90) and blue (90 < pLDDT) representing high and very confidence, respectively, for folded structure. (**C**) PAE matrix, with darker green indicating smaller error, from the AFDB for AF-Q8TD23-F1-v6. (**D**) Left: AFDB prediction scaled and oriented to the same size and direction as its ensemble on the Right: 100 AFX-IDPCG conformations of Zinc finger protein 675 aligned to the C-terminal folded domain colored in grey (residues 231-568), with IDRs in orange and other folded elements from the AFDB template in blue.

For proteins with at least one IDR identified above, Stage 2 classifies the entire protein sequence into one of three categories of increasing structural complexity, which have implications for how conformational sampling for the IDR regions is accomplished in Stage 3. We define the following three categories of proteins containing IDRs as:

1. completely disordered IDPs or proteins with a folded domain and having only N-and/or C-terminal IDRs (often referred to as “tails”);
2. proteins with IDRs between non-interacting folded domains (often referred to as “linkers”), as well as potentially also having N- and C-terminal IDRs; and
3. IDRs within a single folded domain or separating two domains that are likely to form stable interactions (often referred to as “loops”), while also potentially having N- and C-terminal IDRs and/or IDRs between non-interacting folded domains.

Note that the words “tail”, “linker” and “loop” have sometimes been used to imply lack of function beyond non-interacting tethers or elements, but that we use them here only to delineate connectivity, fully appreciating the rich functional repertoires of all IDRs ^37^. Out of the 14,792 identified proteins in the canonical human proteome, 46% (6,763) are in category 1, 22% (3,328) are in category 2, and the final 32% (4,701) are in category 3.

To distinguish categories 2 and 3, we must consider whether the two folded segments connected by an IDR interact (e.g., two parts of a single domain separated by a loop or two tightly bound domains), or whether the folded domains are largely independent of each other (i.e., as “beads-on-a-string”). We use the mean predicted alignment error (PAE) from the AFDB to discriminate between interacting (PAE ≤ 15 Å, category 3) and non-interacting (PAE > 15 Å, category 2) folded domains. Justification for this cutoff is provided by the solute carrier family 26 member 9 (SLC26A9) protein (Supplementary Fig. S2), which has a mean PAE between the folded element of just over 10 Å, a standard PAE cutoff value^38–40^. However, an X-ray crystal structure (RCSB PDB ID 7CH1)^41^ of the SLC26A9 domain aligns to the predicted full-length structure with an RMSD of 1.4 Å, suggesting that PAE values above 10 Å may be found for structurally oriented elements. We therefore took a more conservative approach by increasing the PAE cutoff from 10 to 15 Å. In the AlphaFlex workflow, for proteins that fall into category 3, we generate IDR ensembles while maintaining the AFDB relative orientations for interacting folded domains, whereas IDR ensemble generation for proteins in category 2 does not maintain the AFDB relative orientations for the folded domains.

Stage 3 of the workflow builds the ensembles using information from Stages 1 and 2 and can utilize either IDPConformerGenerator^21,22^ or IDPForge^23^ (Supplementary Fig. S3), both of which provide all-atom ensembles including hydrogen atom positions. IDPConformerGenerator statistically samples backbone torsion angles (ω, φ, ψ) based on distributions for similar sequences in the RCSB PDB, and can also bias the conformations by experimental secondary structure information^21^. IDPForge^23^ is a machine learning tool based on the ESMFold architecture tuned for IDRs based on IDPConformerGenerator^21,22^, CALVADOS^42^ ensembles, and folded proteins from CASP12. IDPForge can generate atomistic conformations of proteins containing folded domains and IDRs, and can generate ensembles under experimental restraints, all at inference time without model retraining. IDPForge ensembles have flexibility in the folded structures at the IDR boundaries^23^, which may be more physically realistic than having fixed backbone coordinates at this boundary, as in the IDPConformerGenerator approach.

There are 3,470 UniProt-annotated^43^ transmembrane proteins in the 14,792 human proteins from AFDB that have predicted IDRs of at least 15 residues. Of these 3,470 membrane proteins, 1,020 contain IDRs that overlap with the membrane-spanning regions as predicted by TMbed^44^ or annotated by UniProt^43^. These may be examples of conditional folding in the presence of a lipid bilayer. Thus, 2,450 transmembrane proteins that contain IDRs were targeted for generating ensembles. For these cases, we applied an additional IDR sampling restraint dictated by an implicit bilayer of 1-palmitoyl-2-oleoyl-sn-glycero-3-phosphocholine (POPC). This uses the experimentally determined average thickness of a POPC bilayer of 39.1 Å^45^ with an adjustable probabilistic interface boundary of 5.15 Å^45^ to allow IDR:head-group interactions and reject IDR conformations that enter the core of the bilayer, as illustrated in Fig. 2.

**Fig. 2.**
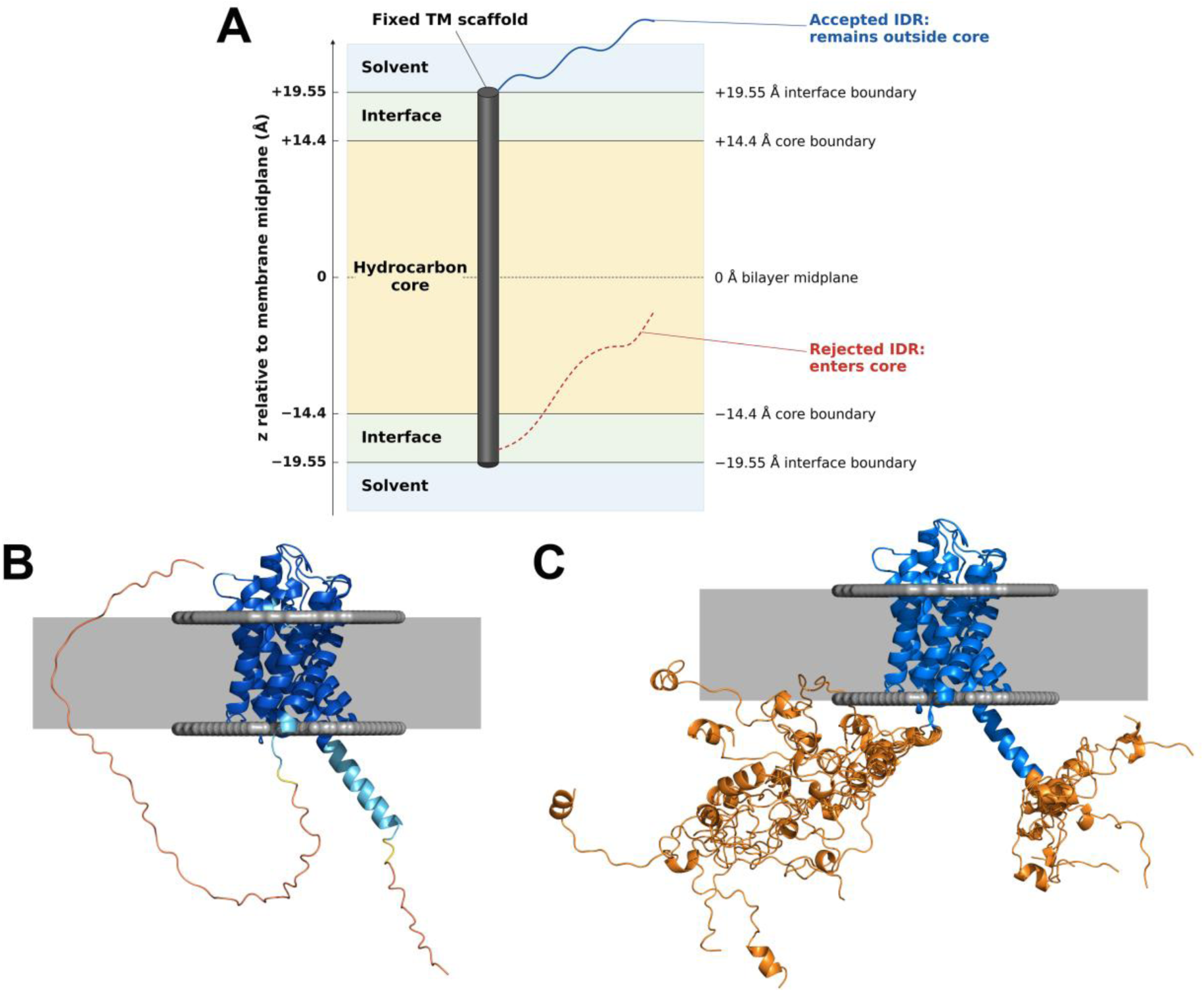
AlphaFlex workflow for transmembrane proteins. (**A**) Schematic of the implicit bilayer to determine allowed and rejected IDR conformations. An average half-thickness of 19.55 Å is used by default^45^. (**B**) AFDB (A0A075B734) prediction of putative aquaporin-7B where the grey slabs are the membrane boundary as predicted by PPM 3.0^67^. The C-terminal low-confidence region can be seen entering the bilayer. (**C**) AFX-IDPCG prediction of putative aquaporin-7B, N = 10 conformations illustrate ensembles with the implicit bilayer method rejecting IDR conformations that enter the hydrocarbon core but allowing some IDR interactions with the implicit bilayer interface. The lighter grey box in (B) and (C) is a schematic of the full extent of the implicit bilayer.

Due to the heavy computational requirements of generating conformations for IDRs between two interacting (fixed) folded domains with IDPConformerGenerator, IDPForge was used to generate category 3 proteins (203 total deposited AFX-IDPForge ensembles). IDPForge is roughly 3 times faster on this task than IDPConformerGenerator. Both IDPConformerGenerator and IDPForge ensembles have been validated to reflect experimental structural properties for IDRs^21,23^. Hence, we provide the community with both sets of ensembles using AlphaFlex; the AlphaFlex ensembles generated with IDPConformerGenerator are referred to as AFX-IDPCG, while those generated with IDPForge are referred to as AFX-IDPForge. The resulting ensembles of 100 conformations per protein have been analyzed for global shape properties, intramolecular contacts, and local secondary structure elements.

At the time of writing this work, over 60% of the targeted proteins from AFDB have AlphaFlex ensembles calculated (a total of 9,315 unique proteins, with 6,763 AFX-IDPCG from category 1, 2,379 AFX-IDPCG from category 2, 203 AFX-IDPForge and 147 redundant and 26 unique AFX-IDPCG from category 3). These have reached Stage 4 and are deposited (or submitted for deposition) into the PED^3^. With the majority of the currently calculated AlphaFlex ensembles being AFX-IDPCG models and evidence for similarity of global structural metrics between the AFX-IDPCG and AFX-IDPForge ensembles (see below), much of the analysis below focuses on AFX-IDPCG models. Note that we use “AlphaFlex” to refer to the general computational workflow, the resulting ensemble models and the collective database.

### Global metrics of ensembles of proteins with IDRs in the absence or presence of folded domains

Previous proteome-wide conformational databases of IDR ensembles were either created by defining an IDR region and then generating ensembles in isolation without fixed endpoints with the folded domains, or by generating full-length proteins including folded domains but with coarse-grained representation, such as in the cases of CALVADOS^24^ and AF-CALVADOS^25^, respectively. Close to all-atom representations exist in the AFDB but are limited to low confidence single-conformer predictions without hydrogen atoms. In contrast, IDPConformerGenerator and IDPForge generate all-atom IDR ensembles in the context of the folded domains^22^ (Fig. 1D). This difference in approach for the AlphaFlex ensembles compared to the AFDB is evident when we analyze global structural metrics of radius of gyration (R_g_), end-to-end distance (R_ee_), hydrodynamic radius (R_h_), solvent accessible surface area (SASA), asphericity (A), and mean curvature (κ).

Fig. 3 presents global structural metrics for AlphaFlex and AFDB full-length IDR-containing protein ensembles/structures as a function of sequence length. While the two sets have some overlapping distributions below sequence length of 1000, AFDB’s hydrodynamic sizes plateau at much lower values (Fig. 3A, B), with R_ee_ being relatively flat as a function of increasing sequence length (Fig. 3B), suggesting that the conformers AFDB generates are anomalously collapsed. In contrast, AlphaFlex shows expected behavior of increasing R_g_, R_h_ and R_ee_ as the length increases (Fig. 3A-C). AFDB has a tendency towards spherical overall shape above a length of 750, while AlphaFlex has a broader distribution of asphericity, particularly for longer sequences, centered around more rod-like shapes (Fig. 3D). SASA follows similar trends between the two sets (Fig. 3E). In addition, we analyzed the curvature for IDRs extracted from AFDB and AlphaFlex. We find that those from AFDB, particularly the longer ones, cluster at lower curvature (∼ 1), reflecting more extended chains, while the curvature of extracted AlphaFlex IDRs have a more normal distribution centered around 2, reflecting the presence of turns in the chains (Fig. 3F).

**Fig. 3.**
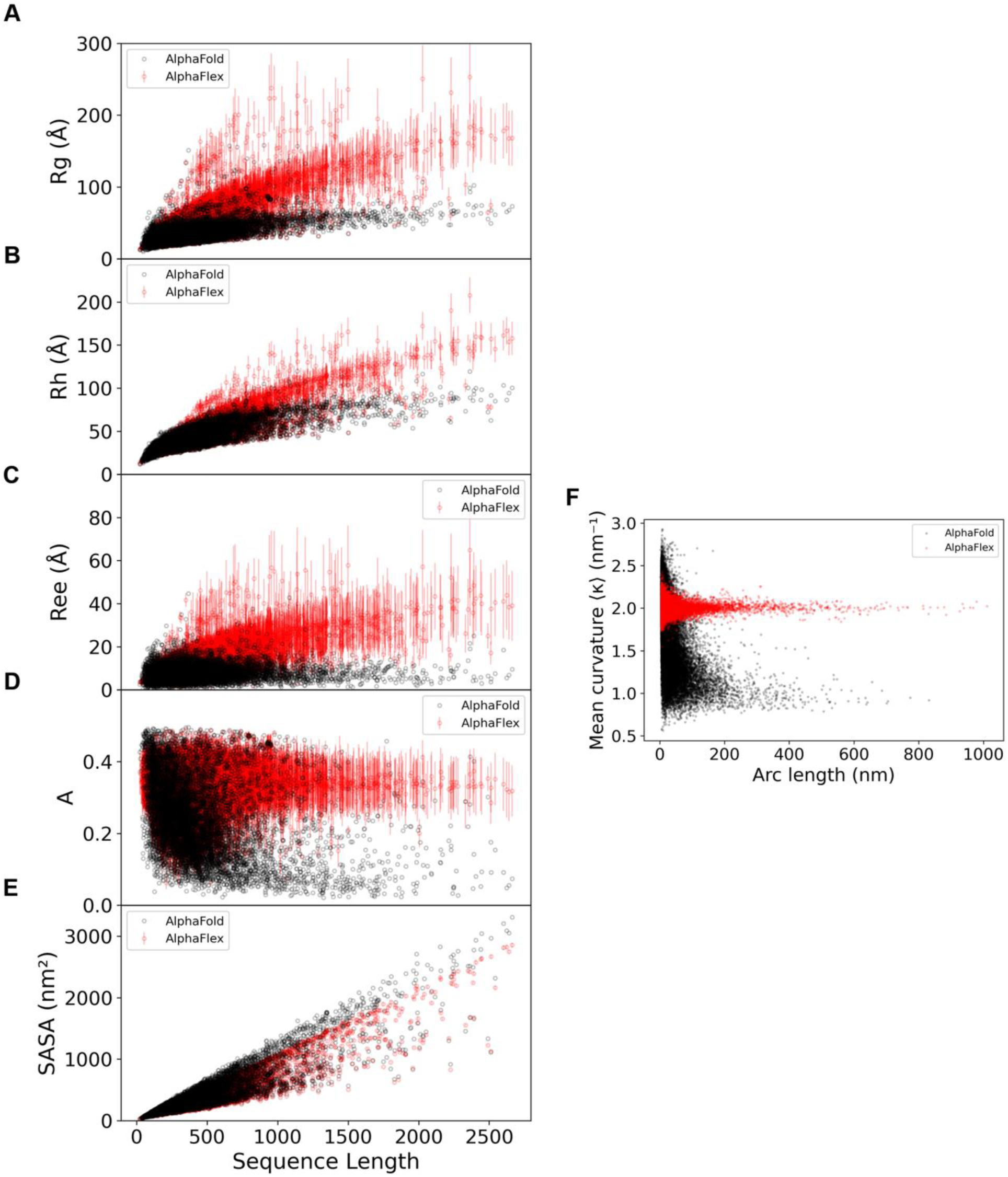
Global structural properties of full-length proteins from AlphaFold database (red) compared to AlphaFlex (black) ensembles and curvature analysis for extracted IDRs. Lines for each red dot represent the mean standard error for the AFX-IDPCG ensembles (N = 100 conformers per protein, total of 9,315 deposited proteins analyzed). Units for A-C are in Angstroms (Å). AlphaFlex plotted in red and AlphaFold plotted in black. (**A**) Radius of gyration (R_g_). (**B**) Hydrodynamic radius (R_h_). (**C**) End-to-end distance (R_ee_). (**D**) Asphericity (A). (**E**) Solvent accessible surface area (SASA). **(F)** Mean curvature (κ) of extracted IDRs versus arc length.

We also compared R_g_, R_h_, R_ee_, A and SASA metrics for isolated IDRs from IDPConformerGenerator and CALVADOS, which performs well in recapitulating experimental hydrodynamic data^46^. Since CALVADOS uses a different definition for IDRs compared to AlphaFlex, only those proteins with the exact same IDR boundaries defined by both CALVADOS and AlphaFlex were analyzed. A total of 1,387 IDR sequences of the human proteome were found to meet this criterion and the normalized Jensen-Shannon divergence metrics of the distribution of structural properties were calculated (Supplementary Table S1.1). IDPConformerGenerator and CALVADOS show no statistically significant differences in 4 of the 5 global structural properties for isolated IDR ensembles (Supplementary Fig. S4), with only small differences found for asphericity (also see Supplementary Table S1.1).

Supplementary Fig. S5 shows analyses of hydrodynamic properties of AlphaFlex (generated with IDPConformerGenerator) and CALVADOS for isolated IDRs compared to the R_g_ and R_h_ values of a null hypothesis for empirical polymer scaling models. Supplementary Fig. S5A,B shows that the concordance correlation coefficient (CCC) values between AlphaFlex and CALVADOS are above 0.96 for both R_g_ and R_h_, indicating close agreement between the two methods, likely reflecting the densely populated smaller IDR lengths, with deviations occurring only between methods only at high values of R_g_ and R_h_. Supplementary Fig. S5C-F shows that the IDPConformerGenerator slope is slightly steeper than the null hypothesis, but all IDR lengths show a linear correspondence, whereas CALVADOS is non-linear and shows outliers for longer sequences. Given the empirical nature of the parameters of the empirical scaling laws, fit a long time ago to sparse data and limited R_g_ and R_h_ ranges, we believe that the experimental and polymer scaling laws support that AlphaFlex and CALVADOS ensembles are physically reasonable.

To understand how the presence of folded domains impacts the global structural properties of IDR ensembles, we analyzed proteins in each of the 3 categories of proteins with IDRs. We find that IDRs derived from the full-length AlphaFlex ensembles versus IDRs generated in isolation by IDPConformerGenerator^21,22^ show no significant differences in all 5 global structural metrics for categories 1 and 2 (Supplementary Fig. S6.1 and S6.2 and Supplementary Table S1.2 and S1.3). However, there are global structural differences for the IDRs derived from full-length ensembles versus those generated in isolation for category 3 proteins. To provide a more straightforward analysis, we consider only the 147 AFX-IDPCG ensembles from category 3 proteins with a single IDR connecting two interacting folded domains (“loop” IDR). Fig. 4 (and Supplementary Fig. S7 and Supplementary Table S1.4) show that IDRs extracted from AlphaFlex ensembles have R_ee_ and asphericity values representing more compact and more spherical conformations than for IDRs built in isolation by IDPConformerGenerator. This is expected given that IDRs between two fixed folded domains predicted to interact with each other would constrain their relative distance and orientation, unlike the IDRs in categories 1 and 2. Comparison of structural property distributions of full-length redundant category 3 proteins between AFX-IDPCG and AFX-IDPForge show no significant differences (Supplementary Fig. S8, Supplementary Table S1.5 and S1.6).

**Fig. 4.**
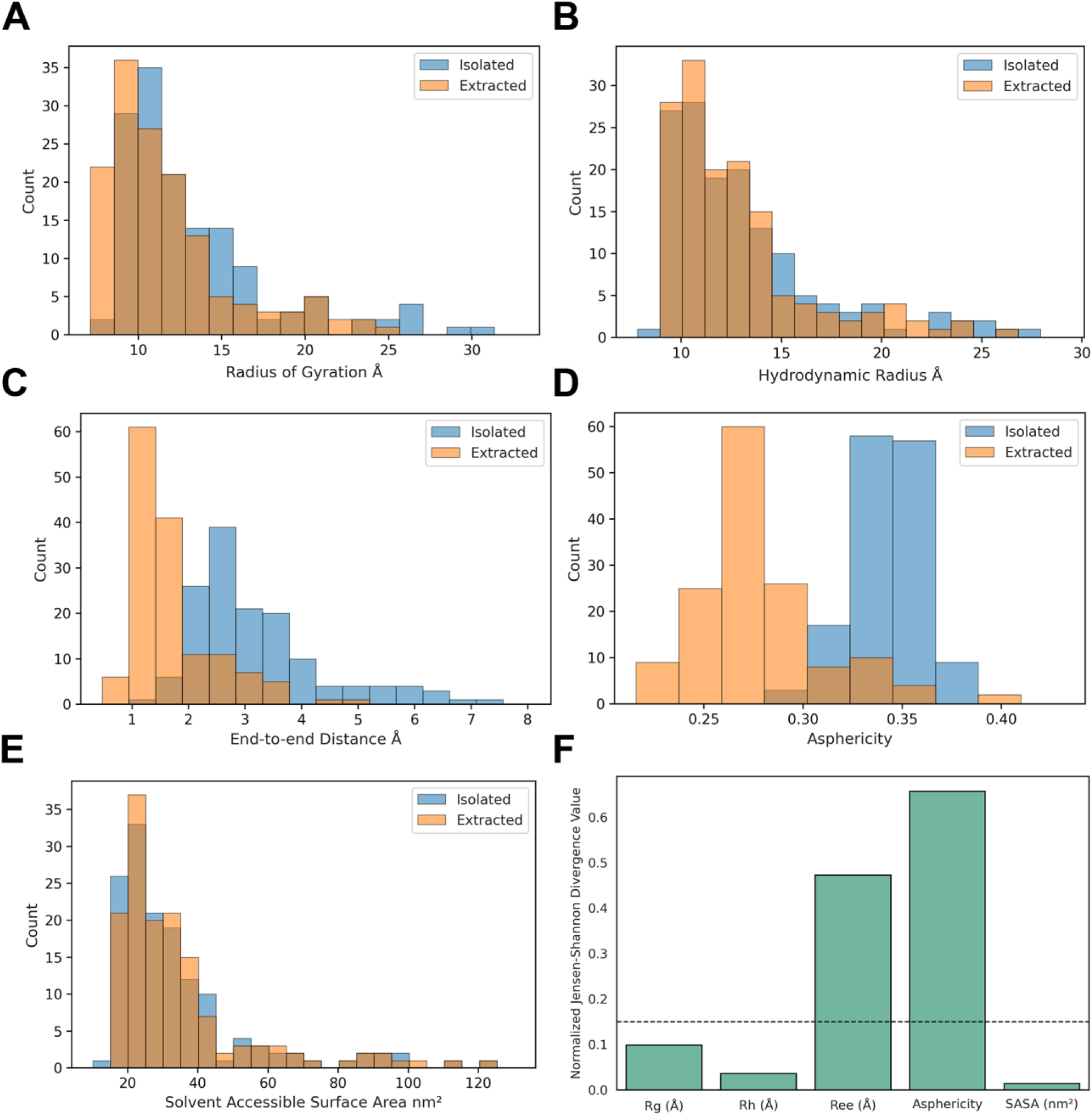
Distributions of ensemble structural properties (category 3, single loop IDR subset, N = 147 proteins) between IDRs generated by IDPConformerGenerator in isolation and IDRs extracted from AFX-IDPCG ensembles. Blue distributions represent IDR ensembles generated in isolation by IDPConformerGenerator. Orange distributions represent IDRs extracted from the context of the folded domain from the AlphaFlex workflow. (**A**) R_g_ (Å) with 1.4 Å bin sizes. (**B**) R_h_ (Å) with 1.1 Å bin sizes. (**C**) R_ee_ (Å) with 0.5 Å bin sizes. (**D**) Asphericity with bin sizes of 0.02 units. (**E**) SASA with 0.5 nm^2^ bin sizes. (**F**) Normalized Jensen-Shannon (JS) statistics of all structural properties. Black dashed line represents a 0.15 cutoff for significantly different properties. SHANK3 is excluded from these analyses since it contains terminal IDRs, IDRs between non-interacting domains and more than one loop IDR between interacting domains, such that it is out-of-distribution with respect to the single loop IDR-containing proteins that currently comprise category 3 proteins.

In order to compare full-length AlphaFlex ensembles with experiment for global characteristics, we found matches for only a limited set of reported proteins with small angle X-Ray scattering (SAXS) data in the SASBDB^47^. We have identified a total of 8 usable SAXS depositions for full-length proteins where the percentage of predicted IDRs is at least 10% of the total number of residues with good experimental quality across q-ranges from 0 to 1.0 A^-1^. We used Pepsi-SAXS v3.0^48^ as a forward-model for SAXS for these 8 IDR-proteins and calculated a normalized χ^2^ value (χ^2^/N) to determine agreement with the SAXS experimental curve (Supplementary Table S2.1). AFDB models significantly deviated from the SAXS experiment, while both AFX-IDPCG and AF-CALVADOS had much better agreement, supporting the expectation of physically meaningful AlphaFlex ensembles with respect to global metrics.

### Local secondary structural features of proteins with IDRs

Many IDRs adopt transient α-helical structure, which is functionally relevant for binding in complexes involving IDRs^5^. Disorder within IDRs is a measure of conformational heterogeneity, not the lack of fractional secondary structure features. Hence, predicted conformational ensembles should not violate the known presence of intramolecular hydrogen-bonds and torsion angle distributions present in the Ramachandran maps of all proteins. We expect the torsion angle sampling for IDRs to fall between distributions found in the full non-redundant high resolution database taken from the RCSB PDB and the torsional distributions of residues not within α-helix or β-strand (defined as loops, L+). We compared AlphaFlex IDRs with AFDB single-state IDRs, CALVADOS IDR ensembles generated in isolation^24^, and AF-CALVADOS IDRs. We used AlphaFlex (N = 9,315 proteins, 100 conformations per IDR), AFDB (N = 14,792 proteins, 1 conformation per IDR), CALVADOS (N_IDR_ = 28,058 isolated IDRs, 100 conformations per IDR), and AF-CALVADOS (19 experimentally benchmarked) ensembles. For the AFDB dataset, IDRs are defined as regions of pLDDT < 70. The entire CALVADOS representation of IDRs for the human proteome was down sampled from 1000 frames to 100 frames and then back-mapped to all-atom using cg2all^49^ to obtain secondary-structure features; the same was done for AF-CALVADOS ensembles. In contrast, IDPConformerGenerator and IDPForge used in the AlphaFlex workflow provide atomistic models directly.

AFX-IDPCG and AFX-IDPForge IDRs show distributions of the α and β regions of the Ramachandran diagram that define the upper and lower bound expected from proteins from the full RCSB and RCSB loops (L+), reflecting unbiased sampling of these energetically favorably basins of torsion angle space in disordered regions^50^. The torsion angle distributions of AFDB low pLDDT regions and CALVADOS IDRs are similar, with reductions in the α region compared to AlphaFlex IDRs and instead favoring the β and coil regions (Table 1). The low level of α region torsion angles within the AFDB low confidence regions has consequences for the chain curvature. As noted previously, longer AFDB IDRs cluster at lower curvature (∼ 1), reflecting more extended chains, while the curvatures of AlphaFlex IDRs cluster around 2 (Fig. 3F), due to the more prominent α region torsion angles.

**Table 1.** Torsion angle distributions of the AFDB, CALVADOS, and AlphaFlex databases. Alpha (α) regions of the Ramachandran diagram are defined as -180° < φ < 10° and -120° < ψ < 45°. Beta (β) regions of the Ramachandran diagram are defined as -180° < φ < 0° and -180° < ψ < -120° or 45° < ψ < 180°. Coil regions are defined as φ, ψ torsion angles not belonging to α or β regions. Torsion angle distributions of the entire IDPConformerGenerator database which include RCSB PDB X-Ray crystal structures with resolution better than or equal to 2.0 Å as of 2024 are given as a reference, including for all residues and those not within regular α-helical or β structure (L+). Protein regions with pLDDT < 70 from the AFDB were analyzed (N = 14,792 structures). 100 conformers for each of the CALVADOS ensembles (N = 28,085 IDRs, back mapped to all-atom), AFX-IDPCG extracted IDRs (N = 17,071 extracted IDRs from 9,315 proteins), and AFX-IDPForge extracted IDRs (N = 203 ensembles, one IDR per ensemble) were analyzed. Values for the AlphaFlex and CALVADOS databases are given as the mean percentage ± standard deviation for all structural conformations. Values with an * do not have an associated mean standard deviation since they are not ensembles.

| **Method** | **% $\alpha \pm STD$** | **% $\beta \pm STD$** | **% coil $\pm STD$** |
| --- | --- | --- | --- |
| *RCSB PDB Database* | 50.1* | 43.5* | 6.4* |
| *RCSB PDB Database L+* | 35.8* | 53.1* | 11.1* |
| *AFDB* | 17.6* | 58.9* | 23.5* |
| *CALVADOS* | 21.1 ± 1.0 | 57.9 ± 1.9 | 21.0 ± 1.5 |
| *AFX-IDPCG* | 46.9 ± 7.4 | 46.8 ± 7.0 | 6.3 ± 3.0 |
| *AFX-IDPForge* | 36.6 ± 5.3 | 44.9 ± 7.7 | 18.5 ± 4.7 |

We also calculated the percentages of DSSP^51^-defined secondary structure of IDRs from AlphaFlex, AFDB, CALVADOS and AF-CALVADOS (Supplementary Table S3). While the degree of stable hydrogen-bonded helix in IDRs is not precisely known, the experimental observation of fractional α-helix^52,53^ and biological importance of α-helical elements in IDR recognition elements^54,55^ suggests that it should be above 10%. We observe that extracted IDRs from the AlphaFlex ensembles have the highest percentage of DSSP-defined α-helices, with a uniform 23% to 25% for IDRs across all sizes, including those with at least 1000 residues (Supplementary Table S3.3), consistent with IDRs having substantial fractional α-helical structure^52,53^. Although AFDB has ∼10% DSSP-defined α-helix across all the pLDDT < 70 residues, this decreases to ∼6% for IDRs with sequence lengths above 100, and only ∼1% for IDRs with sequence length of 1000 amino acids (Supplementary Table S3.4). CALVADOS completely lacks hydrogen-bonded helix in its IDR ensembles, which is expected since coarse grain protein models represent residues as single beads which fail to coordinate hydrogen-bonding and electrostatic interactions required to form local secondary structures. AF-CALVADOS IDR ensembles have a mean DSSP-defined helical population of ∼8% (Supplementary Table S3.1), likely due to use of AlphaFlex IDR boundaries allowing some folded structure to be included in the analysis.

In order to compare full-length AlphaFlex ensembles with experiment for local features, we searched databases for NMR chemical shift data in the BMRB^56^. We identified a total of 13 NMR chemical shift depositions for full-length proteins where the percentage of predicted IDRs is at least 10% of the total number of residues. We used UCBShift v2.0^57^ as a forward-model for NMR chemical shift data, and only the chemical shifts for the IDRs were assessed (Supplementary Table S2.2). Although there are only a small number of NMR data comparisons, AFDB models were found to deviate more beyond experimental uncertainties than either AFX-IDPCG or AF-CALVADOS, and supports the expectation of physically meaningful AlphaFlex ensembles.

### Intramolecular domain contacts of proteins with IDRs

AlphaFlex ensembles also have different intramolecular contacts between IDRs and folded domains than AFDB. The PAE metric can be used to quantify the likelihood of contacts between IDRs (D) and folded domains (F). The mean PAE values 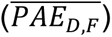 reveal that approximately 6.7% (or 992) of the 14,792 AFDB proteins contain at least one 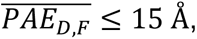 for IDR and folded domain regions separated by at least 5 amino acid residues to remove trivial local contacts. Due to the dynamic nature of IDRs, these intramolecular IDR:folded-domain interactions would be expected to sample a distribution of intramolecular Cα (IDR) to Cα (folded domain) distances. Fig. 5 highlights two examples, the putative CENPB DNA-binding domain-containing protein 1 (UniProt ID B2RD01, Fig. 5A-C) and Antigen peptide transporter 1 (UniProt ID Q03518, Fig. 5D-F). This comparison shows AFDB representations having close IDR:folded-domain contacts (Fig. 5A,D) with AlphaFlex exhibiting intramolecular IDR:folded-domain contacts that vary from close to distant (Fig. 5B,E). The distribution of distances is biologically relevant, providing accessibility to folding domains and IDR segments containing post-translational modification or protein binding sites, while also providing potential regulatory contacts.

**Fig. 5.**
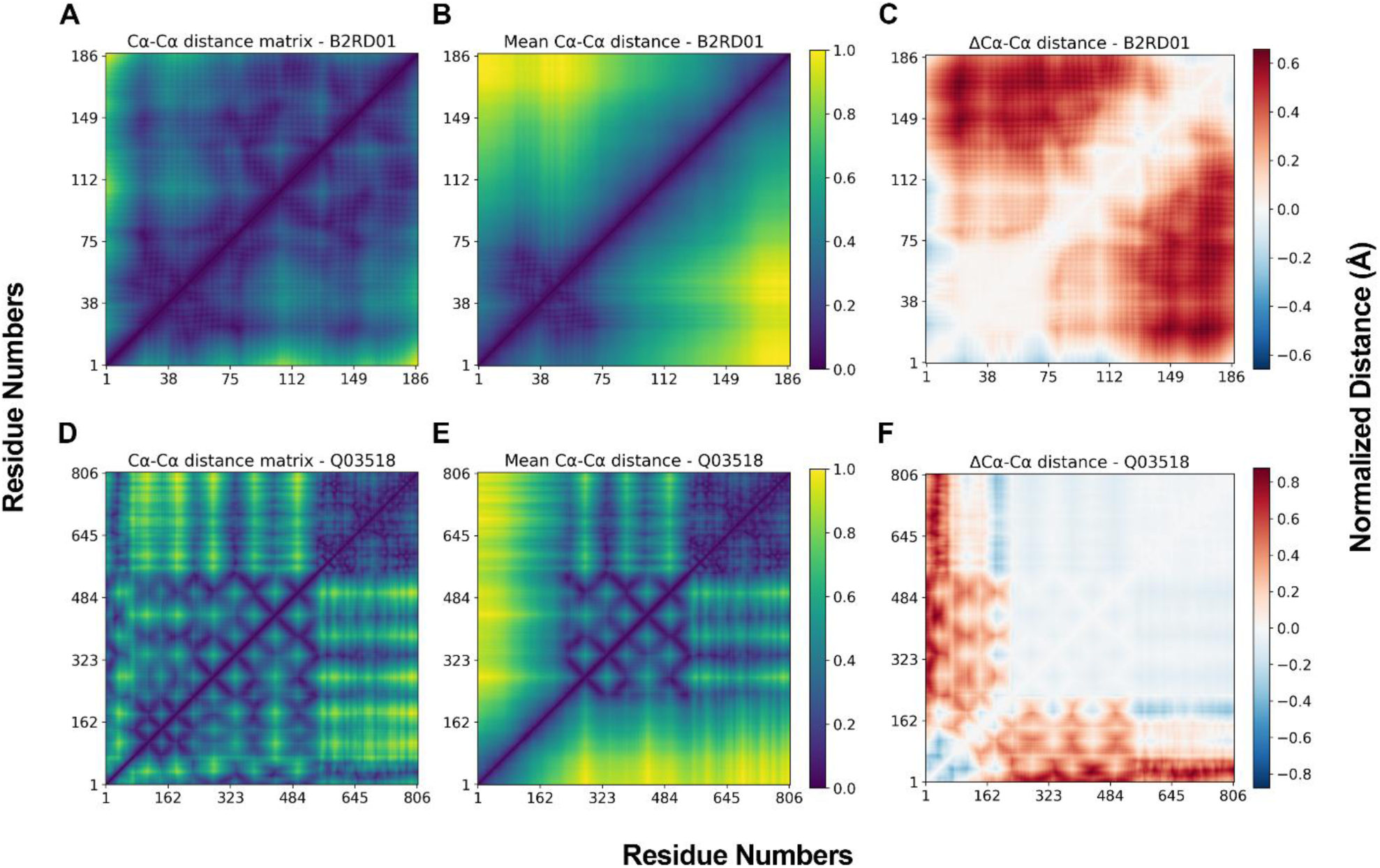
Comparison of Cα-Cα distance matrices derived from AFDB and AlphaFlex representations. First column depicts the Cα-Cα distance matrix from AFDB^2^ predicted structure. The second column depicts the mean Cα-Cα distance matrix from N = 100 AlphaFlex conformers. Matrices have been normalized (with 1 representing the longest distance). The last column depicts the differences of normalized Cα-Cα distance matrices where AFDB is subtracted from mean AlphaFlex Cα-Cα distance matrices. (**A-C**) Category 1 Putative CENPB DNA-binding domain-containing protein 1 (UniProt ID B2RD01). Total protein length: 187 aa. IDR ranges: 1-21, 66-187. (**D-F**) Category 2 Antigen peptide transporter 1 (UniProt ID Q03518). Total protein length: 808 aa. IDR ranges: 1-114, 135-231.

These Cα-Cα distance matrices also reflect the different IDR boundaries between AFDB (based on pLDDT confidence values) and AlphaFlex, the latter having more residues defined as IDRs. The AlphaFlex ensembles for Putative CENPB DNA-binding domain-containing protein 1 (B2RD01) have N-terminal (residues 1-21) and C-terminal (residues 66-187) IDRs while AFDB has confident prediction of structure for residues 84-140. The AlphaFlex ensemble for Antigen peptide transporter 1 (Q03518) contains an N-terminal (residues 1-114) IDR as well as a linker IDR (residues 135-231) and ends with a C-terminal folded domain, while AFDB has three confidently predicted folded regions (residues 89-92, 115-134, and 232-808). Misrepresenting an IDR as a potential folded domain is not only a mistake in assignment but also under-represents the conformational heterogeneity of IDRs which have the potential to be closer to or further from the folded domain. In addition, IDRs between two folded domains that do not have tight interactions 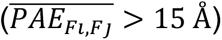 lead to wide ranges of distances between folded domains (*Fi* and *Fj*) as shown in Fig. 5C and 4F.

Given the importance of electrostatic interactions in IDPs, IDRs, and condensates, there is a clear expectation that folded domain chemistry influences IDR interactions through electrostatics. We note that IDPConformerGenerator and IDPForge only implicitly represent these interactions through extraction of short structural fragments from the PDB for IDPConformerGenerator and, for IDPForge, through training data on IDPConformerGenerator, CALVADOS, and CASP12 folded proteins where physically motivated interactions are represented. Hence, although IDPConformerGenerator and IDPForge do not explicitly account for energetic terms, we have tested this hypothesis with a chimera experiment involving a protein with multiple acidic and basic patches on the folded domain, and an IDR enriched with positive, negative, or polar-neutral residues through its sequence. Given that IDPConformerGenerator and IDPForge are expected to behave similarly, we do this computational experiment with IDPConformerGenerator.

We chose an ankyrin domain family protein (UniProt ID A6NI47) as a test case since it has clear basic and acidic surface-accessible patches distant from the IDR:folded-domain junction point (Supplementary Table S4). The chimeric test cases had 4 different 132 aa N-terminal IDRs with a unique sequence repeat to test different IDR chemistries, while the C-terminal IDR was kept as the wild-type sequence. The positive electrostatic sequence case contained repeated “KKRKKRSGPGKRSK” sequences, the negative electrostatic case contained repeated “EDEDEDSGPGDESE” sequences, and the polar-neutral case contained repeated “STNQGSGTQNSPGS” sequences, in addition to the WT sequence. Supplementary Fig. S9.1 shows that the AFX-IDPCG chimeric ensembles had overall preferences for favorable electrostatic surface interactions in which IDRs interactions with acidic and basic surface patches of the folded domain dominate the ensemble; anecdotal evidence for these IDR-folded domain pairings are seen in Supplementary Fig. S9.2. Overall, these results provide evidence for AlphaFlex ensembles containing physically meaningful conformations between folded domains and disordered regions.

### Biological relevance of AlphaFlex IDR ensembles

The AlphaFlex ensembles across all three categories are structurally different than the AFDB representations. This difference is not merely a physical manifestation of possible conformations but is critically relevant for biological function. Single-state structural representations found in AFDB can inhibit understanding of biological mechanisms by their often-misleading implications, while ensemble representations, such as those provided by AlphaFlex, can illuminate regulatory roles of IDR-containing proteins, including scaffolding and other binding processes^10,30^. Although AlphaFold2 can generate multiple models given a protein sequence, the publicly accessible ColabFold (AlphaFold2 Google Colab)^58^ was only able to generate 5 conformations using paid compute credits for sequences longer than 1000 aa. The 5-conformation ensembles generated by AlphaFold2 do not sample a wide distribution of R_g_ values, unlike AlphaFlex conformations (Supplementary Fig. S10.1). Ensemble representations sampling diverse hydrodynamic properties are especially critical for proteins in categories 2 and 3 with IDRs between non-interacting and interacting folded domains. In the case of category 2 proteins, in which the IDRs are generated between non-interacting folded domains (in addition to any N- and C-terminal IDRs), the AlphaFlex ensembles demonstrate a distribution of R_g_ and R_ee_ consistent with a “bead-on-a-string” picture, whereas the AFDB provides a single compact conformation of the full-length structure with IDRs appearing as lower curvature chains that avoid steric clashes, often appearing as IDR “lassos” around central folded domains (Fig. 6 A-F). The distributions of R_g_ and R_ee_ show that AlphaFlex ensembles have on average a broader distribution for compaction and extension. The difference in scale when viewing all structures of the ensemble at once is apparent in Fig. 6, where a single AFDB structure is about 1/5^th^ the scale of the full ensemble sampled by AlphaFlex. This would have significant implications for the “reach” of the protein, with the AlphaFlex conformations enabling the IDR chain to bridge farther distances important for scaffolding.

**Fig. 6.**
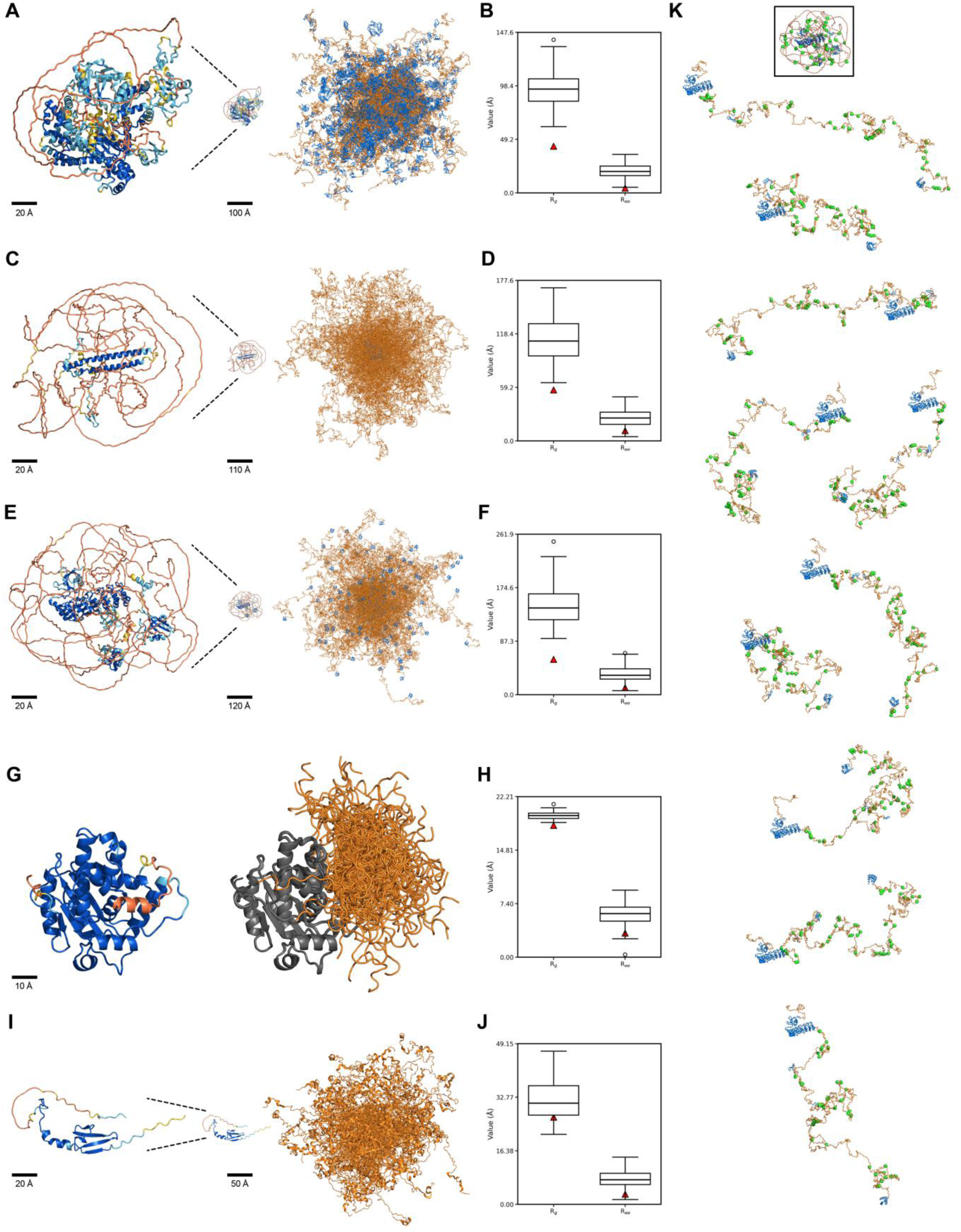
Comparisons between AlphaFold database predictions and AlphaFlex ensemble representations and biological relevance for five example proteins. (**A-B**) Zinc finger MYM-type protein 6 (UniProt ID O95789), 1325 aa long, 449 disordered residues. Ensembles are presented aligned to the C-terminal folded domain (residues 808-1325) colored in grey. (**C-D**) Histone deacetylase complex subunit SAP130 (UniProt ID Q9H0E3), 1048 aa long, 918 disordered residues. Ensembles are presented aligned to the C-terminal folded domain (residues 916-1033) colored in grey. (**E-F**) SH3 and multiple ankyrin repeat domains protein 3 (SHANK3, UniProt ID Q9BYB0), 1806 aa long, 1360 disordered residues. Ensembles are presented aligned to the N-terminal folded domain (residues 76-414) colored in grey. (**G-H**) CCR4-NOT transcription complex subunit 7 (CNOT7, UniProt ID Q9UIV1), 285 aa long, 25 disordered residues. Ensembles presented aligned to the N-terminal folded domain (residues 1-260). (**I-J**) Eukaryotic translation initiation factor 4E-binding protein 2 (4E-BP2, Uniprot ID Q13542), 120 aa long fully predicted to be disordered. The first column depicts AFDB representations of the full-length protein colored by pLDDT ranges (see Fig. 1 legend). The second column depicts AlphaFlex ensemble representations of the full-length protein where orange regions are defined to be disordered from the union of 5 disorder metrics with blue regions being the resulting defined folded domains with structures taken from AlphaFold. AFDB structures are also presented in the second column if a different scale is applied to better represent the scale of the conformational ensemble. The third column are boxplots of the R_g_ and R_ee_ (Å) for the AlphaFlex ensemble with the single R_g_ and R_ee_ values of the AFDB structure overlaid (red triangle). (**K**) AlphaFlex conformers (N = 10) of SHANK3 highlighting residues that undergo post-translational modifications (defined by UniProt) as green spheres. Topmost structure is the AFDB model of SHANK3 outlined in a black box, colored by AFDB pLDDT (defined in Fig. 1 legend). Bottom 10 conformations are the same scale and orientation. IDRs are colored orange while folded domains are blue for the AlphaFlex conformers.

In addition, Fig. 6 demonstrates that the hydrodynamic properties (R_g_ and R_ee_) have clear functional consequences, highlighting folded domains with binding interactions required for function (blue folded domains in Fig. 6 A-F) and post-translational modification sites (Fig. 6, green residues) that are accessible in the AlphaFlex ensembles but more occluded in the AFDB model. Zinc finger MYM-type protein 6 is a nuclear protein involved in regulating cell morphology^28^, with 8 DNA- (and potentially RNA-) binding zinc fingers and many post-translational modifications (PTMs), including SUMOylation, ubiquitination and phosphorylation, providing binding sites for SUMO interacting and ubiquitin interacting domains^43^. The AFDB model with the IDRs wrapped around the folded zinc finger domains does not provide accessibility for nucleic acid binding or the enzymes needed for PTMs and does not have space for the SUMO and ubiquitin additions (also see Supplementary Fig. S10.2). In contrast, the AlphaFlex ensemble with its range of conformations provides accessibility and space for binding partners, enzymes and PTMs (Fig. 6A-B).

Another case is the histone deacetylase complex subunit SAP130 which functions in the assembly, regulatory interactions, and/or enzymatic activity of the mSin3A corepressor complex that plays a role in transcription^29^. Similar to the Zinc finger MYM-type protein 6, SAP130 is highly post-translationally modified, including arginine methylation, phosphorylation, SUMOylation, and ubiquitination^43^ (Fig. 6C-D and also Supplementary Fig. S10.3). The AFDB model with very long IDRs wrapped around a small, folded domain does not provide accessibility to the folded element that engages in regulatory interactions within the mSin3A complex or for the enzymes needed for PTMs and subsequent regulatory binding for SAP130, in contrast to the AlphaFlex ensemble that has a range of accessibilities.

An important example of a scaffold protein is the SH3 and multiple ankyrin repeat domains protein 3 (SHANK3, Fig. 6E-F, K), which phase separates and mediates assembly of the postsynaptic density (PSD), a biomolecular condensate on the post-synaptic side of neuronal synapses that controls biological responses to synaptic stimulation, including local actin cytoskeletal dynamics^30^. SHANK3 has multiple folded binding domains (ankyrin repeats, SH3, PDZ, and SAM) and long IDRs that are required for multivalent interactions leading to phase separation and scaffolding functions. In addition, it has many PTM sites^43^, highlighted in Fig. 6K. As in the previous examples, the AFDB representation puts all the folded binding domains in the center of the long IDRs, which wrap around in a spherical shape, incompatible with the phase separation and scaffold function of the protein and accessibility of enzymes to the PTM sites. The AlphaFlex model, in contrast, demonstrates the long reach of the individual binding domains in this primarily “beads-on-a-string” scaffold.

Other examples highlighting the biological relevance of AlphaFlex ensembles relative to AFDB structures include enzymes and conditionally folding IDRs. CNOT7 is a subunit in the major cellular polyA-RNA deadenylase CCR4-NOT complex that regulates RNA stability^59^, with a low confidence α-helix predicted by the AFDB model to lie in the active site, which would block enzymatic function, while the AlphaFlex conformational ensemble provides heterogeneous conformations, most with the active site open, consistent with enzymatic activity, with some closer to the active site that could be important for regulating activity (Fig. 6G-H). The IDP 4E-BP2 has a conditionally folding IDR. In isolation, NMR data shows a fractionally populated α-helix stabilized by binding to eIF4E (the mRNA 5’ “cap-binding protein”)^60^, which inhibits mRNA translation by blocking eIF4E binding to eIF4G in the translation initiation complex^61^. Translation is stimulated by multi-site phosphorylation of 4E-BP2^62^, leading to conditional folding of a β-sheet domain that includes the residues that are fractionally α-helical in isolation, reducing its binding to eIF4E and enabling eIF4G to compete^26^. The AFDB 4E-BP2 conformation has the β-sheet domain, along with some α-helix, confusing the mechanism of phosphoregulation of translation by presenting a folded conformation that lacks accessibility to the key phosphoregulatory sites and that cannot bind to eIF4E. In contrast, the AlphaFlex ensemble (Fig. 6I-J) represents 4E-BP2 as an IDP with up to 50% fractional α-helical structure for the eIF4E-binding α-helix and accessibility to the phosphorylation sites (Supplementary Fig. S11), consistent with its functional state in regulating translation. An example of the biological intuition provided by AlphaFlex is for the transmembrane protein aquaporin-7B protein (UniProt ID A0A075B734). While the C-terminal IDR of the protein is not well-characterized experimentally, cytoplasmic IDRs of membrane proteins are critical for regulatory interactions^19^. The AFDB conformation places the C-terminal IDR in the core of the bilayer, while the AlphaFlex ensemble shows it to be outside of the bilayer (Fig. 2), accessible for regulatory functions.

## DISCUSSION

Physically meaningful models of full-length proteins are a key first step to generating biological insight regarding structure-function relationships and are better described as structural ensemble-function relationships. These ensemble representations are essential, since single conformations, including those provided by the AlphaFold database, do not explain most biology. AFDB incorrectly models confident structures for conditionally folding IDRs^16^, with particular challenges for those that have multiple conformations in different biological contexts, such as 4E-BP2^26^. More generally, the single conformation AFDB provides for IDRs is often unphysically collapsed into “lasso-like” conformations surrounding clustered folded domains that only satisfy steric clash restraints rather than physically reasonable global shape properties. Such structural models also imply contacts between domains that are not confidently predicted by AFDB’s own PAE metric, but just as importantly they (unintentionally) lead to structures that violate known biological principles and functional roles. Thus, while the AFDB has been transparent that their IDR structure predictions are of low confidence, they are qualitatively wrong in their physical dimensional scale and, most critically, they are biologically implausible.

The AlphaFlex ensembles provide powerful insights into the functional roles of the proteins with intrinsic disorder by more effective modeling of conformational sampling. AlphaFlex utilizes the strength of AlphaFold^2^ predictions of folded domains together with IDPConformerGenerator and IDPForge, low-computational-cost (Supplementary Table S5) atomistic IDR modeling strategies that are well-validated against experimental solution data^21,23^ for proteins with intrinsic disorder. Both show strong agreement with not just experimental R_g_, but NOE and PRE distance data, as well as chemical shifts. Furthermore, AFX-IDPCG ensembles have good agreement with full-length SAXS data and better match for NMR chemical shift data compared to AFDB structures. Additionally, AlphaFlex IDRs sample more local secondary structure compared to AF-CALVADOS ensembles, predominantly fractional α-helical elements, due to the use of IDPConformerGenerator and IDPForge^21,23^. The observed local structure of IDRs within AlphaFlex ensembles leads to greater curvature than low confidence regions of AFDB, which is also much better aligned with function, particularly for helical binding elements.

Our AlphaFlex workflow enables interchangeable modeling strategies and holistic approaches to define IDR boundaries on a proteome-wide scale, and also provides an accessible method to easily visualize full-length ensembles containing IDRs through the PED^3^. Other recently described methods with different philosophies for interpreting IDR regions include Ensemblify^38^ and AF-CALVADOS^25^. Both Ensemblify and AF-CALVADOS heavily rely on AlphaFold’s pLDDT and PAE metrics to define IDR boundaries and interacting folded domains. Although we agree that the PAE score is a valuable metric for interactions between confidently predicted folded domains, the per-residue pLDDT is not the best predictor of intrinsic protein disorder^63^, which can lead to biologically inconsistent IDR assignments that obscure functional states, particularly in the case of conditionally folding IDRs^63^. Thus, AlphaFlex defines IDRs based on the union of various bone-fide disorder predictors.

Using these boundaries and PAE values, our AlphaFlex workflow discriminated between three categories of proteins with IDRs, based on presence of N- and/or C-terminal IDRs, IDRs between non-interacting domains, and IDRs between stably interacting domains. Ensembles for all categories are created in the context of the folded domains to obtain representative structures that provide better models for understanding biological function, for example mediating complex formation or cellular scaffolding^5,10,30^. For a large subset (2,450) of transmembrane proteins, we also providing ensembles of these proteins built in the presence of the implicit bilayer. We found that folded domains bias the structural landscape of IDRs in the context of full-length proteins compared to the same IDRs generated in isolation for folded domains that interact (category 3). However, although terminal IDRs (category 1) or linker IDRs between non-interacting folded domains (category 2) exhibit no differences in global structural properties when compared to IDRs generated in isolation, full-length ensembles must be modeled to avoid steric clashes and generate R_g_ and R_ee_ distributions that enable the reach of folded domains and accessibility to fulfill scaffolding and regulatory binding functions, such as in large multi-domain complexes.

Additionally, AlphaFlex ensembles can delineate between favorable and unfavorable IDR:folded-domain intramolecular interactions. Chimeric IDR sequences in the ankyrin domain family protein with net-negatively and net-positively charged N-terminal IDRs have large populations of favorable contacts with different acidic and basic surface-accessible patches on the folded domain. These observations are consistent with a growing view that IDR:folded-domain intramolecular contacts are likely to be common in full-length multidomain proteins, even when they are weak, transient, and not apparent from global ensemble properties alone. Because IDRs are covalently tethered to folded domains, the effective local concentration of potential interaction partners is high, allowing charge-complementary, hydrophobic, aromatic/π, and hydrogen-bonding contacts to become populated as dynamic intramolecular states. Such interactions can bias IDR ensemble dimensions, tune accessibility of linear motifs and post-translational modification sites, mediate autoinhibition or allosteric regulation, and influence partner binding or phase-separation behavior^37^. Recent experimental and computational studies support this framework by showing that IDR ensemble biases can persist in cells and can be reshaped by tethered folded domains^64^, while emerging predictors and biophysical studies demonstrate chemically specific interactions between IDRs and folded-domain surfaces^65^. Therefore, modeling IDRs in the context of their native folded domains, as done in AlphaFlex, is important not only for avoiding steric artifacts but also for identifying weak intramolecular interactions that may be functionally relevant.

For the AFDB entries of the human proteome having IDRs of 15 residues or more (N = 14,792), we have currently calculated AFX-IDPCG ensembles for all category 1 proteins, AFX-IDPCG ensembles for a subset of proteins (71%) in category 2, a small subset (4%) of AFX-IDPForge and AFX-IDPCG ensembles for the difficult category 3 set, and 231 transmembrane proteins built with the restriction of an implicit bilayer. Complete sets for both AFX-IDPCG and AFX-IDPForge across all three categories are in progress. Current and future ensembles generated by the AlphaFlex workflow may be used as information-dense machine learning training data. Since IDPConformerGenerator enables custom secondary structure sampling (CSSS) biased by NMR chemical shifts^21^, experimental data may be used to regenerate AFX-IDPCG ensembles where available. In addition, AlphaFlex may be used to model protein ensembles beyond those of the canonical human proteome, leveraging the strengths of experimentally validated protein prediction and modeling strategies.

Having AlphaFlex ensembles deposited in the PED is a community resource that permits more proteome-wide analyses and biological interpretations of the full protein conformational space, highlighting the importance of global and local structural properties of IDRs in the context of full-length proteins for function. Furthermore, UniProt currently links to PED ensembles (listed just below AlphaFold in the “Structure” section), making accessibility of our conformational ensembles of proteins with IDRs extremely straightforward; for example, see Epsin-2 (UniProt ID O95208) and Homeobox protein Hox-C10 (UniProt ID Q9NYD6). At the time of writing, AlphaFlex ensembles have already been useful for scientists as initial models for MD simulations, lowering the barrier for modeling full-length proteins containing IDRs^66^. Importantly, AlphaFlex predictions provide a physically relevant conformational ensemble of monomeric proteins in dilute solution as well as in the context of a membrane bilayer for transmembrane proteins. Future efforts include modeling the remainder of human proteins containing IDRs, modeling the proteome’s other isoforms and conformational ensembles in other environments, including dynamic complexes (homo- and hetero-multimers) and biomolecular condensates, representing the next pressing frontiers.

## ACKNOWLEDGEMENTS

Z.H.L. and J.D.F.-K. acknowledge Dr. Simon Sharpe and Dr. Hue Sun Chan for their computational hardware contributions of AlphaFlex ensemble calculations. S.C.E.T and A.M.Monzon acknowledge ELIXIR, the research infrastructure for life-science data; views and opinions expressed are however those of the author(s) only and do not necessarily reflect those of the European Union or the European Research Executive Agency. The authors acknowledge Dr. Jonathon Ditlev for helpful discussions and scientific input.

## METHODS

### Creating the AlphaFlex database of IDRs

AFDB structures and protein FASTA sequences of the canonical human proteome (Proteome ID UP000005640) were downloaded from the AlphaFold^2^ and UniProt^43^ databases, respectively, on July 24^th^, 2024. Four well-established predictors of intrinsic disorder, metapredict^33^, flDPnn^34^, ADOPT^35^, and SPOT-Disorder^36^, were applied to the protein sequences, with a cutoff of 15-consecutive amino acids predicted to be disordered used to identify IDRs. In addition to IDP/IDR predictors, we also included as an indicator of disorder, the AFDB pLDDT confidence metric being below or equal to 70 as AFDB documentation recommends disregarding unconfident predicted structures^2^. Since some IDR prediction tools have sequence length limitations, we maximized coverage across the human proteome by taking the union of the IDR boundaries from all 5 indicators of intrinsic disorder to maximize coverage of IDRs.

### Generating conformer ensembles using IDPConformerGenerator

IDPConformerGenerator builds IDRs by sampling backbone torsion angles (ω, φ, ψ) based on a Monte Carlo statistical approach for distributions of similar sequences (in fragments of 1 to 5 amino acids) within a databased containing non-redundant X-ray crystal structures with a resolution ≤ 2.0 Å from the RCSB PDB^21^. IDRs can be generated in isolation or in the context of an existing set of coordinates, as described previously^22^. After generating IDRs in the context of the folded domains from AFDB models, filtered by the IDR definitions above, PDBFixer is used through OpenMM 8^68^ to add hydrogens to the sidechains of conformations generated by FASPR^69^, the default sidechain packing protocol within IDPConformerGenerator^21,22^. After protonation, an implicit solvent, fixed charge, energy-minimization molecular dynamics simulation was performed using the *AMBER99sb*^70^ forcefield through OpenMM^68^ for a maximum of 2 ns while fixing the backbone atoms to resolve any sidechain steric clashes. Finally, 100 conformers were collated into a single file using pdb-tools^71^. Specific protocols for directing IDPConformerGenerator to generate category 1, 2, and 3 protein ensembles can be found in the AlphaFlex subdirectory in the IDPConformerGenerator GitHub repository (<u>github.com/julie-forman-kay-lab/IDPConformerGenerator</u>). Computational hardware resources used to calculate the AlphaFlex ensembles can be found in Supplementary Table S5.

### Generating conformer ensembles using IDPForge

IDPForge^23^ is a denoising diffusion probabilistic model (DDPM) designed to generate atomistic structural ensembles for both fully disordered proteins and disordered regions within the context of folded domains. The model adapts the triangular attention, structure module, and core architectural components of the protein structure prediction method ESMFold^72^ to learn the sequence-dependent and inter-residue spatial relationship ultimately refining conformations iteratively starting from noise. To predict ensembles that contain terminal or linker IDRs, IDPForge takes a disordered residue range and a folded template as input and uses a mixed timestep sampling strategy that only denoises the disordered segments while conditioning on the sequence and residue features of the folded structure. Distinctively, IDPForge ensembles capture fractional secondary structure in disordered regions while making a smooth transition into the folded domains. Further details can be found in the paper^23^.

Modifications to the IDPForge method to enable calculations of category 1, 2, and 3 protein cases can be found in a subdirectory within the IDPForge GitHub repository (<u>github.com/THGLab/IDPForge</u>). All IDPForge calculations were run in a Conda environment on the Savio HPC cluster, with template generation performed primarily on CPUs and conformer generation/relaxation completed using NVIDIA A40 GPUs (Supplementary Table S5).

### Categorizing transmembrane proteins and creation of the implicit bilayer

The UniProt REST-API was used to identify transmembrane and intramembrane annotations to obtain 5,229 transmembrane proteins from the AFDB human proteome. These transmembrane proteins were cross-referenced with the AlphaFlex IDR database to retain only UniProt IDs of transmembrane proteins with at least 1 IDR. TMbed^44^ was then used on the 3,470 transmembrane proteins with IDRs to predict transmembrane topology/boundaries. The TMbed boundaries were cross-referenced again with the AlphaFlex IDR database to retain only cases where the IDR does not overlap the membrane spanning regions predicted by TMbed, resulting in 2,450 proteins targeted for ensemble generation.

Building of transmembrane proteins utilized the same philosophy of the other AlphaFlex protein cases but include an implicit POPC-derived bilayer with thickness dimensions from experimental data (hydrocarbon core of 14.4 Å, and headgroup at 19.55 Å)^45^ infinitely spanning the XY plane at Z = 0, with a Z-thickness of ± 19.55. The TM Z-axis at Z = 0 is estimated by taking the median Z-coordinate position of Cα atoms from predicted TM segments. The folded template is then rotated and centered at Z = 0 so implicit bilayer slab boundaries can be applied. IDRs that penetrate the hydrocarbon core are rejected while those that have interface contacts get exponentially penalized as more IDR atoms approach the hydrocarbon core. The rejection logic checks the number (N) of IDR atoms’ Z-coordinate (*z_abs,i_*) at the interface levels (*d_i_)* where 0 is near solvent facing and 1 is near the core boundary and applies an adjustable interface rejection strength of *s = 0.15*. The formula for calculating the probability of accepting an IDR (*P_accept_*) is

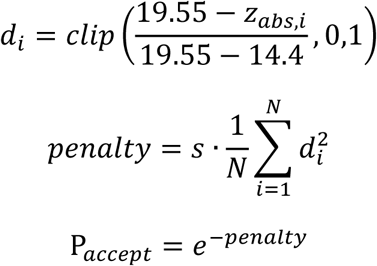

In the example of Fig. 2, manually processing the AFDB structure with PyMOL^73^ and building the bilayer boundaries with PPM v3.0^67^ took 30 minutes (web-tools), while generating N = 100 conformations with the AlphaFlex implicit bilayer method took 45 minutes using 50 CPU workers.

### Analysis of global and local ensemble structural properties

A Python script integrating MDTraj^74^, NumPy^75^, and HullRad^76^ was used to calculate structural and sequence features. Structural properties include radius of gyration (R_g_), hydrodynamic radius (R_h_), end-to-end distance (R_ee_), asphericity (A), and accessible surface area (SASA). HullRad^76^ was used to calculate R_h_. The *compute_rg* function in MDTraj^74^ was used to calculate R_g_. The *shrake_rupley* function in MDTraj^74^ with a probe radius of 0.14 Å was used to calculate SASA. R_ee_ was calculated by finding the Euclidean distance between the first and last residues using MDTraj^74^. Similarly, Cα-Cα distances were calculated using MDTraj^74^ by filtering the coordinates for “CA” atoms and obtaining a Euclidean distance matrix from the subset of coordinates. Asphericity was calculated by computing the eigenvalues (*λ*) from the inertia tensor and taking the 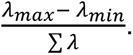 Backbone torsion angles (ω, φ, ψ) were calculated using the IDPConformerGenerator *torsion* module. Fractional secondary structure was calculated by using DSSP^51^ through the IDPConformerGenerator^21^ *sscalc* module. IDPConformerGenerator groups DSSP categories into L, H, and E, where L consists of DSSP hydrogen bonded turn (T), bend (S), loops and irregular elements (NULL), residues in isolated β-bridge (B), π-helix (I), and poly-proline type II helices (P). H consists of α-helix (H) and 3_10_ helices (G). Lastly E consists of extended β-strands that participates in hydrogen-bonded β-sheets.

Local curvature and arc length were calculated using the Cα coordinates of each conformation. The backbone was treated as a discrete piecewise-linear curve comprising of atom position *r_i_* where *i* is the total number of residues in the IDR. Computation of the local curvature *κ*_i_ (Å^-1^) used two distinct vectors between 3 residues *r_i-1_, r_i_,* and *r_i+1_* and the angle *θ_i_* between them as follows^77^.

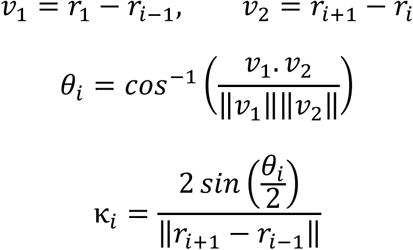

The arc-length of a protein chain segment was computed as the sum of consecutive Cα-Cα distances as follows.

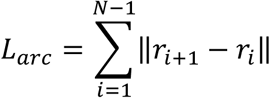

We have reported these values for ensembles processed at the time of writing in the supplemental materials archive. The Python scripts for structural analysis are also available in the supplemental materials archive. Visualization of structural conformations and ensembles were performed in PyMOL v3^73^; secondary-structure elements may not be correctly visualized by PyMOL due to the nature of multi-state conformations.

IDPConformerGenerator and CALVADOS IDRome hydrodynamic properties were also compared against polymer scaling models. For R_g_ scaling^78^ we used *R_g_ = 2.54 x N^0.522^* and an R_h_ scaling law^79^ of *R_h_ = 2.49 x N^0.509^* as seen in Fig. S5.

### Benchmarking against experimental SAXS and NMR chemical shift data

Both UniProt IDs and the primary sequence were used to scan the SASBDB^47^ and BMRB^56^ to find matching depositions for full length proteins. NMR NOE and PRE data were also considered when searching through the BMRB, however no full-length protein matches were found. The curated 100-protein NMR spectra dataset was also considered however no full-length matches for proteins containing IDRs were found^80^. 22 SASBDB entries were found, however, only 8 were usable since the rest were experimental data of the protein captured in a complex or had unusable I(q) errors, described below. All 13 matching NMR chemical shift depositions were used for benchmarking.

A Python script was used to perform independent quality control of experimental SAXS data based on intensity uncertainties. For each experimental SAXS.dat file, (q), I(q), and σ(q) were parsed. We quantified noise using relative error (σ/|I|) and signal-to-noise ratio (|I|/σ), with special emphasis on the early screening window (q[0.10–0.20]); curves were rejected if this region already showed poor quality (default thresholds: median (σ/|I| > 0.25), 75th percentile (> 0.50), (> 20%) of points with (σ/|I| > 0.50), median SNR (< 4), or (> 2%) non-positive intensities). To avoid reacting to isolated spikes, we applied rolling-median smoothing and identified the first sustained run of “bad” points ((σ/|I| > 0.50) or SNR (< 2), at least 8 consecutive points). Curves with no sustained deterioration passed; curves that became noisy only in the final 20% of the screened (q)-range were retained as tail-noisy; curves with deterioration before the tail were flagged for cropping at the last reliable (q); and curves failing early-region criteria were rejected.

### Surface chemistries of folded domains and IDR:folded-domain interactions

To annotate the surface chemistry of folded domains, solvent-accessible surface area (SASA) was calculated at the residue level, and relative solvent accessibility (RSA) was defined as

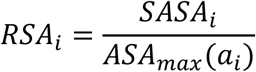

where *SASA_i_* is the solvent-accessible surface area of residue *i,* and *a_i_* is the amino acid identity, and *ASA_max_(a_i_)* is the theoretical amino-acid specific solvent accessibility^81^. Residues were classified as surface-exposed if *RSA_i_ ≥ 0.25*.

The chemistry of each folded-domain patch was summarized using net charge, net charge per residue (NCPR), fraction of charged residues (FCR), positive and negative residue fractions, hydrophobic residue fraction, and aromatic/π-residue fraction. Residue charges were assigned as

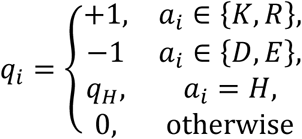

where *q_H_ = 0.1* by default. Hydrophobic residues were defined as {A, V, I, L, M, F, W, Y, P, C}. π-capable residues were defined as {F, Y, W, H, R}, where F, Y, and W contribute to aromatic ring centroids, H contributes to the imidazole centroid, and R contributes the guanidinium centroid.

For each conformer in the ensemble, residue-level contacts were calculated between IDR and the folded-domain residues. Contacts were computed using distance-based cutoffs between atoms or residue-level centers. A residue pair was counted at most once for a given interaction class, even if multiple atom-atom contacts occurred between the same two residues. However, a single IDR residue could contribute multiple residue-pair contacts if it contacted multiple folded-domain residues. When multiple IDRs were present, the IDR set for determining contacts could be restricted to all IDRs, only the N-terminal IDR, only the C-terminal IDR, or user-specified IDR intervals. Folded-domain residues were defined as all residues outside the full set of annotated IDR intervals, whereas the IDR residues for defining contacts were drawn only from the selected target interval(s). Unless otherwise specified, contacts made by residues immediately adjacent to the IDR:folded-domain junction were excluded using an anchor-distance cutoff of 10 residues. Thus, an IDR residue *i* was included in the set for defining contacts only if *d*_anchor_(*i*) ≥ *d*_cut_, where *d*_anchor_(*i*) is the sequence distance from residue *i* to the selected IDR:folded-domain boundary and *d*_cut_ = 10 residues by default. This filter was used to eliminate trivial tether-proximal contacts.

Charge contacts were calculated between oppositely or similarly charged residue pairs using a 6.0 Å cutoff. Opposite-charge and like-charge residue-pair counts were denoted as *N_opp_* and *N_like_* respectively. Opposite-charge pairs satisfy *q_i_q_j_ < 0,* whereas like-charge pairs satisfy *q_i_q_j_ > 0.* Hydrophobic contacts were defined as side-chain carbon or sulfur atoms from hydrophobic residues within 4.5 Å, giving a residue-pair count *N_hydrophobic_*. Cation-π contacts were defined between residue-level cation centers and aromatic/imidazole π-system centroids within 6.0 Å, evaluated in both IDR-cation/folded-domain-π and folded-domain-cation/IDR-π orientations. Cation centers were defined as the Lys terminal ammonium atom *NZ*, the Arg guanidinium centroid from atoms *NE*, *CZ*, *NH1*, and *NH2*, and the His imidazole centroid when His was assigned a positive effective charge. The aromatic/imidazole acceptor set for cation-π contacts was {F, Y, W, H}. these corresponding pairs was denoted *N_cationπ_*. π–π contacts were defined between side-chain π-system centroids within 6.0 Å. The π-system set was {F, Y, W, H, R}, where F, Y, and W were represented by aromatic ring centroids, H by the imidazole centroid, and R by the guanidinium centroid. Due to Arg-Arg pairs having like-charge repulsion but also contributing to a favorable π-π contact, Arg-Arg contacts were still favorable but rewarded less than other π–π contacts to account for charge repulsion. The resulting residue-pair count was denoted *N_ππ_.* Hydrogen-bond-like contacts were calculated using donor/acceptor heavy atoms. A hydrogen bond was counted when the donor (*D*)–acceptor (*A*) distance (*d*), hydrogen (*H*)–acceptor distance, and donor–hydrogen–acceptor angle satisfied

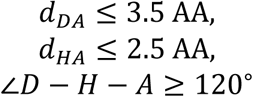

For each residue pair *i, j* and contact type *t*, ensemble contact occupancy was calculated as

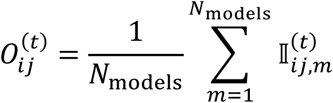

Where 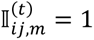 if residue pair *i, j* formed contact type *t* in model *m*, and 0 otherwise.

Interaction classes were not treated as mutually exclusive. Therefore, a single IDR:folded-domain residue pair could contribute to multiple contact classes when multiple criteria were satisfied; for example, Arg-His could contribute to charge, cation-π, π-π, and hydrogen-bond-like contact counts depending on geometry.

### Statistical comparison of distributions of ensemble structural properties

A Python script using NumPy^75^ and other default packages were used for the statistical comparison of different ensemble properties. The Jensen-Shannon (JS) statistic was chosen for this study because it is symmetric and remains finite even when two histograms place data in disjoint bins^82^. To calculate JS, the Kullback-Leibler (KL) divergence (or relative entropy) must be obtained to measure information loss incurred when distribution *q* is used to approximate distribution *p* and vice versa. Since KL is asymmetric and unbounded, *KL(q||p)* and *KL(p||q)* are combined symmetrically inside the JS divergence metric. The formula to calculate the normalized JS statistics is given below; An ε = 10^-12^ value was added to every bin since any zero entry in one histogram may cause a term to become undefined. 10^-12^ is much smaller than any real probability mass after normalization.

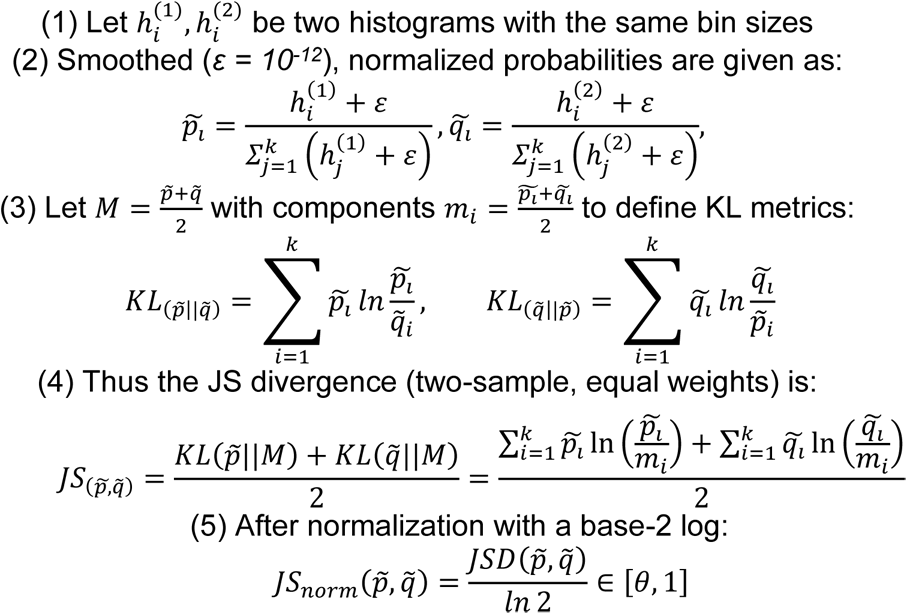

### Validation of AlphaFlex Ensembles

AlphaFlex ensembles after generation underwent a 4-step validation process. First, multi-conformation ensemble PDB files were checked to ensure there were exactly 100 unique conformations per ensemble, by ensuring there were 100 non-redundant REMARK MODEL lines after the ensemble was compiled with pdb-tools^71^. Secondly, the full-length protein sequence in the ensemble was checked to match the full-length sequence from the AFDB template by converting all the residue 3-letter-codes to 1-letter and performing a simple string equivalency check. Third, stereochemistry of the amino acids in the IDRs was double checked to ensure that only L-enantiomers were generated. This involved assuring that the backbone torsion angles of non-glycine residues did not sample inaccessible regions of the Ramachandran plot (i.e., torsion angles were not found in the 0° < φ < 180° and 0° < ψ < -180° region of the plot). Ramachandran plots for each ensemble are also viewable within the PED, where torsion angles that slightly deviate from the expected distribution are highlighted in red. Lastly, text-encoding was double checked to be “utf-8” and any files found to be encoded in “utf-8-sig”, “cp1252”, and “latin-1” using the “decode” function were re-encoded to “utf-8” by using a simple Python “with” statement, specifying “encoding=’utf-8’”.

### Extracting conformer ensembles from CALVADOS and AF-CALVADOS representations

To ensure that the number of conformations from the CALVADOS ensembles match the number of conformations from the AlphaFlex ensembles, every 10^th^ frame of the CALVADOS ensembles were extracted using MDTraj^74^. Since CALVADOS provides coarse grain representations, cg2all^49^ was used to recapture atomistic information. We note that cg2all is computationally intensive for converting proteome scale datasets for downstream analysis, and this step took 28 CPU days on a single AMD Threadripper 2990wx.

### Automated processing of IDR ensembles within the Protein Ensemble Database

PED ensembles for each unique UniProt ID, including AFX-IDPCG and AFX-IDPForge ensembles, are contained within the same PED ID, as distinct sub-ensembles. PED uses a deposition web service (https://deposition.proteinensemble.org/)^3^ to manage data associated with protein ensembles. This service supports metadata handling, computation and visualization of structural features, and automated quality assessment through a standardized validation pipeline. While the deposition interface was originally designed for manual submissions, where users upload ensemble files and accompanying metadata, here we leveraged the underlying API endpoints to enable automated, high-throughput deposition.

In this work, ensemble deposition was executed on the HPC infrastructure at the University of Padova (BioComputing UP Lab). Job submission and scheduling were managed through DRMAAtic^83^, which provides a RESTful interface to HPC resources via the Distributed Resource Management Application API (DRMAA). This integration allowed secure, efficient, and scalable deposition workflows, enabling bulk submission of ensembles without manual intervention.

## Data and materials availability

The AlphaFlex workflow and analysis scripts are available under a subdirectory of the IDPConformerGenerator repository on GitHub after update v0.8.X (github.com/julie-forman-kay-lab/IDPConformerGenerator). The code for conformer generation with IDPForge is available at https://github.com/THGLab/IDPForge.git Ensembles calculated in this study have been uploaded to Zenodo (10.5281/zenodo.17684897), with all AlphaFlex ensembles being uploaded to the Protein Ensemble Database in a rolling fashion. Users can search the PED with the keyword “AlphaFlex” to identify AFX-IDPCG and AFX-IDPForge ensembles.

