## Supplementary Information for "AlphaFlex: Ensembles of the human proteome representing disordered regions"

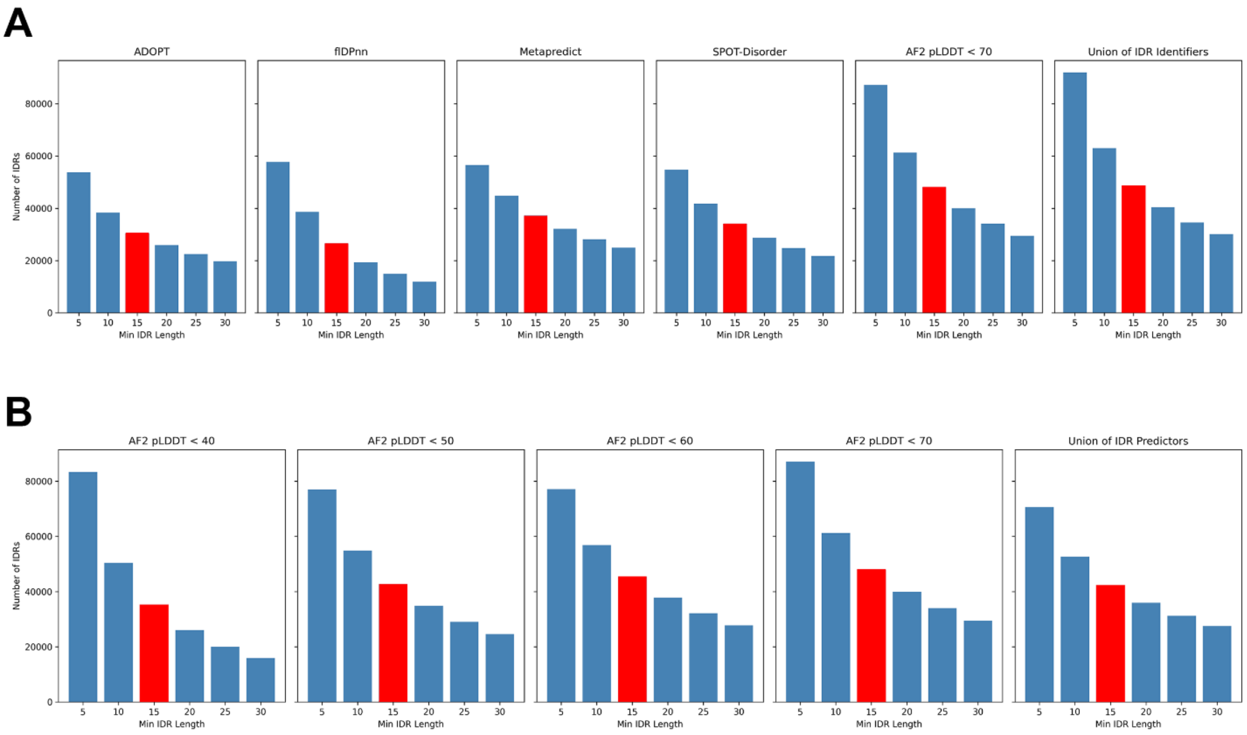


**Supplemental Figure S1. Numbers of IDRs defined using different cutoffs for minimum consecutive disordered residues and using different metrics for disorder.** Red bars represent the minimum length cutoff (15 consecutive residues) chosen for this study. (**A**) Different predictors of intrinsically disordered regions from left-to-right: ADOPT, flDPnn, metapredict v3, SPOT-Disorder, AlphaFold pLDDT < 70, and the union of all 5 indicators of protein disorder. (**B**) Numbers of IDRs defined solely based on AlphaFold pLDDT values (<40, <50, <60, and <70) and defined solely on the union of disorder predictors (i.e., not including pLDDT < 70).

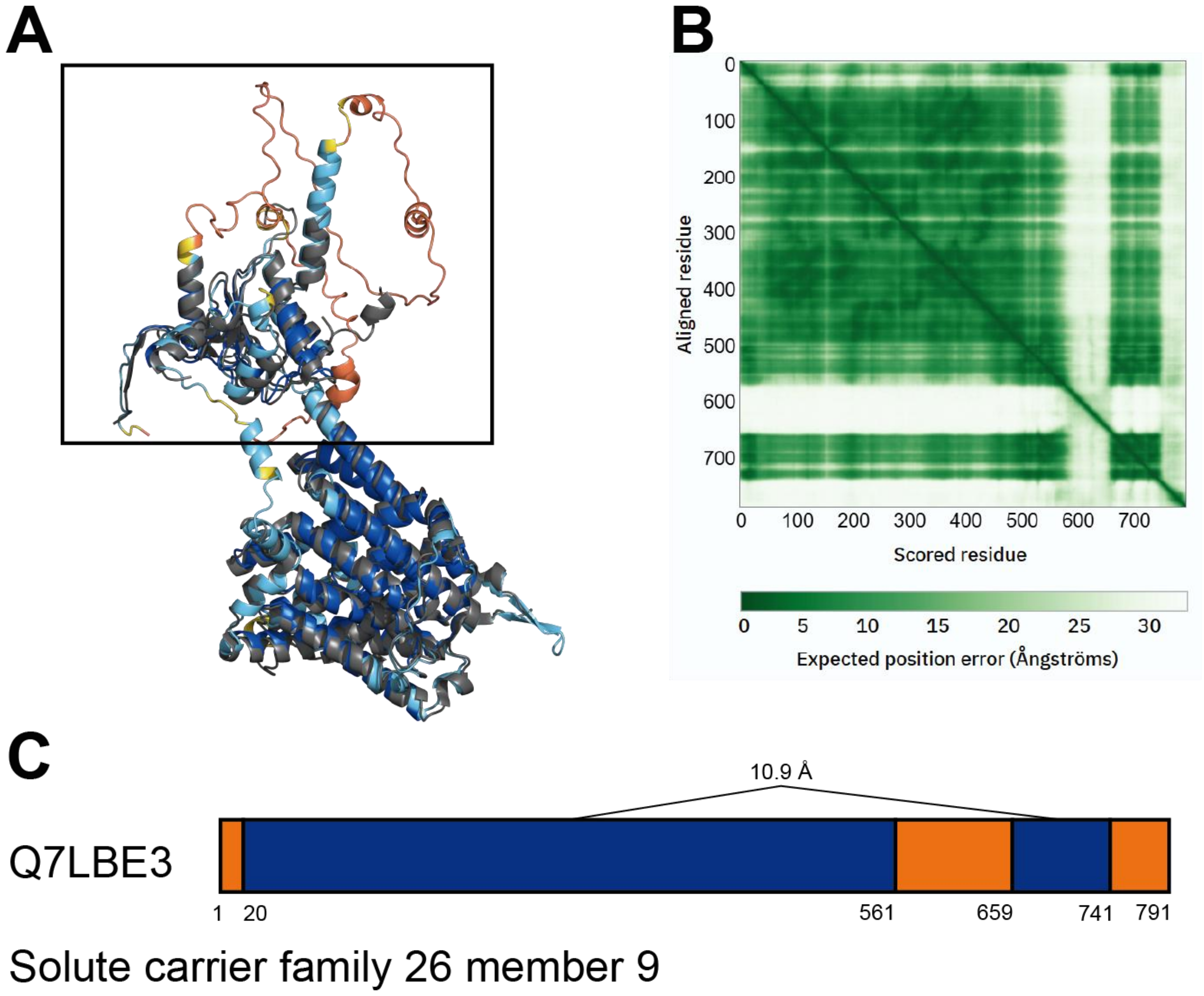


**Supplemental Figure S2. Solute carrier family 26 member 9 (SLC26A9, UniProt ID Q7LBE3) example from the AlphaFlex database compared to experimentally solved structure (RCSB PDB ID 7CH1).** (**A**) The predicted and experimentally solved structure of SLC26A9 have been aligned together and visualized using PyMOL[1]. The AFDB model is colored by its pLDDT score as defined in the AFDB with yellow and orange representing low (50 < pLDDT < 70) and very low (pLDDT < 50) confidence, respectively, and light blue (70 < pLDDT < 90) and blue (90 < pLDDT) representing high and very confidence, respectively[2]. The grey structure represents the experimentally solved chain A of SLC26A9 (PDB ID 7CH1)[3]. The black box highlights the folded domains on either side of the second IDR (residues 562-659). (**B**) The PAE matrix of SLC26A9 obtained from the AFDB. (**C**) Protein regions in orange are predicted to be disordered. Protein regions in blue by extension are the defined folded domains. The $\overline{PAE_{i,j}}$ for a pair of folded domains *(i)* and *(j)* connected by solid lines are given.

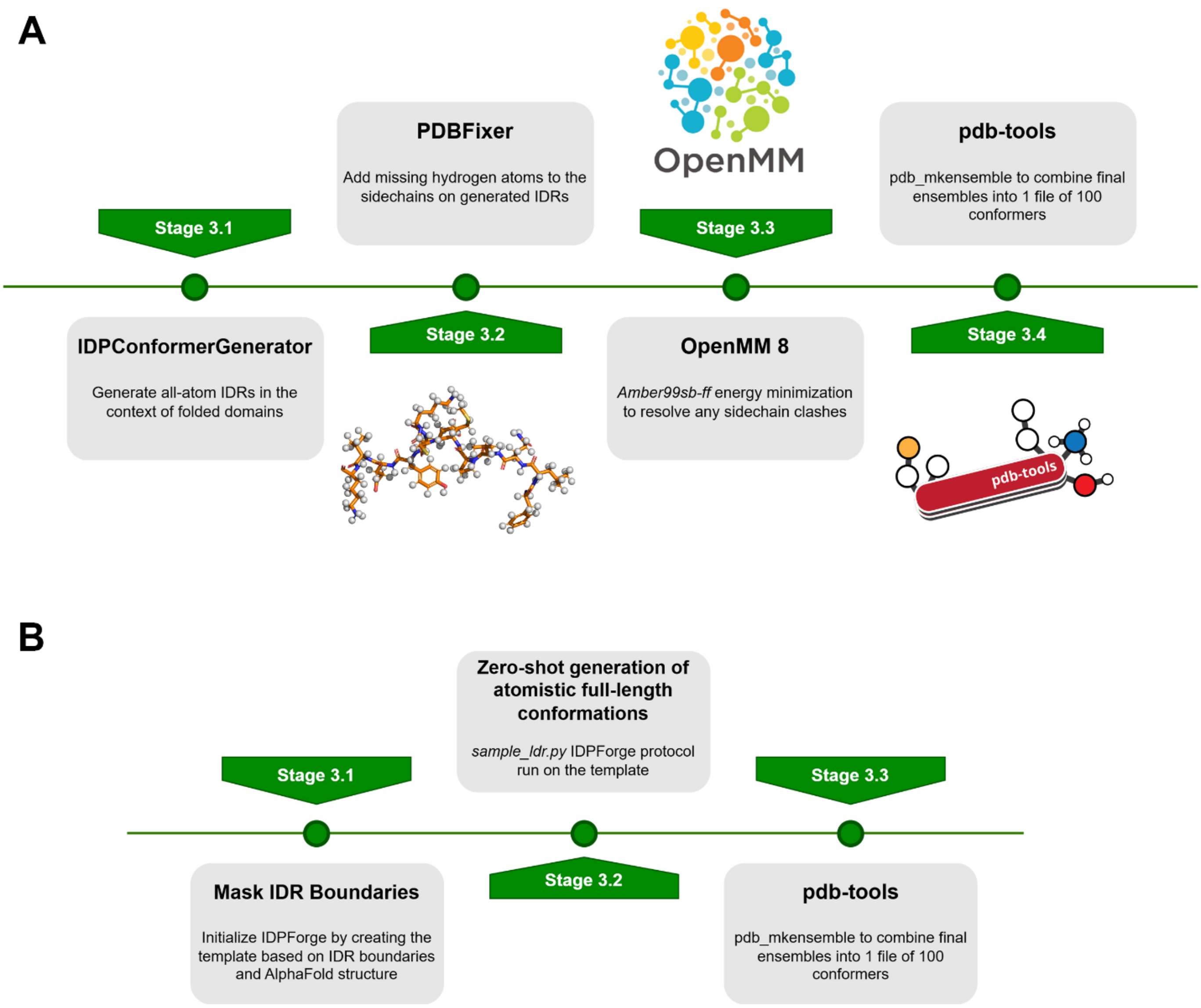


**Supplemental Figure S3. Detailed workflow for AFX-IDPConformerGenerator and AFX-IDPForge pipelines.** (**A**) IDPConformerGenerator workflow variation of Stage 3 from Fig. 1. (**B**) IDPForge workflow variation of Stage 3 from Fig. 1.

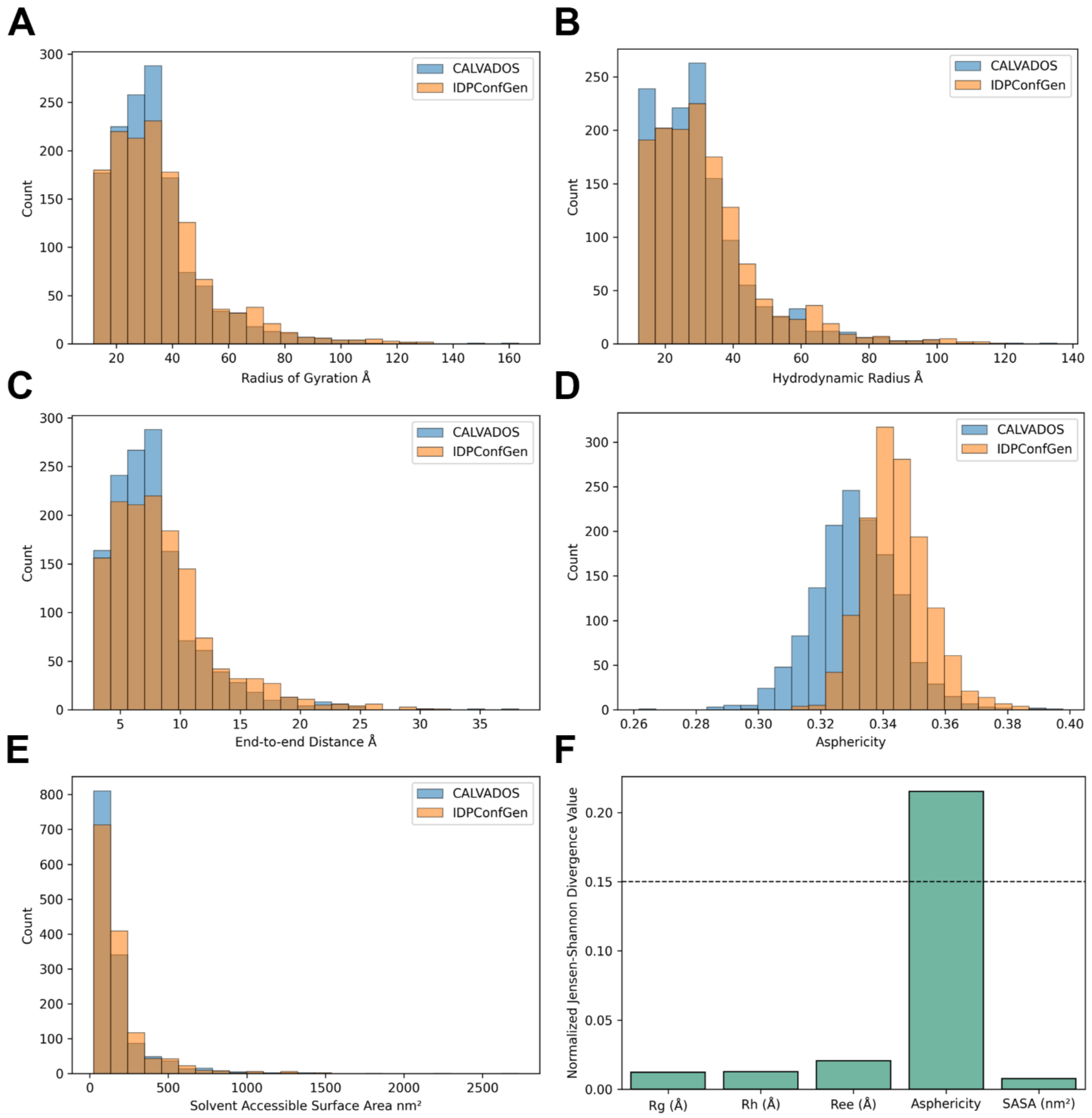


**Supplemental Figure S4. Distributions of ensemble structural properties for 1,387 matching IDR sequences between CALVADOS and IDPConformerGenerator ensembles when generated in isolation.** Blue distributions represent CALVADOS ensembles. Orange distributions represent IDPConformerGenerator ensembles. IDR sequences were derived from the AlphaFlex definition. (**A**) $R_g$ (Å) with 1 Å bin sizes. (**B**) $R_h$ (Å) with 1 Å bin sizes. (**C**) $R_{ee}$ (Å) with 1 Å bin sizes. (**D**) Asphericity with bin sizes of 0.01 units. (**E**) SASA with 20 $nm^2$ bin sizes. (**F**) The normalized Jensen-Shannon divergence value ($JS \in [0, 1]$) with a cutoff of 0.15 for substantial differences indicate that the only structural property that was statistically significant between the CALVADOS and IDPConformerGenerator distributions was asphericity, with IDPConformerGenerator ensembles on average more spherical than CALVADOS ensembles. No statistical differences between the distributions for $R_g$, $R_h$, $R_{ee}$, and SASA are found.

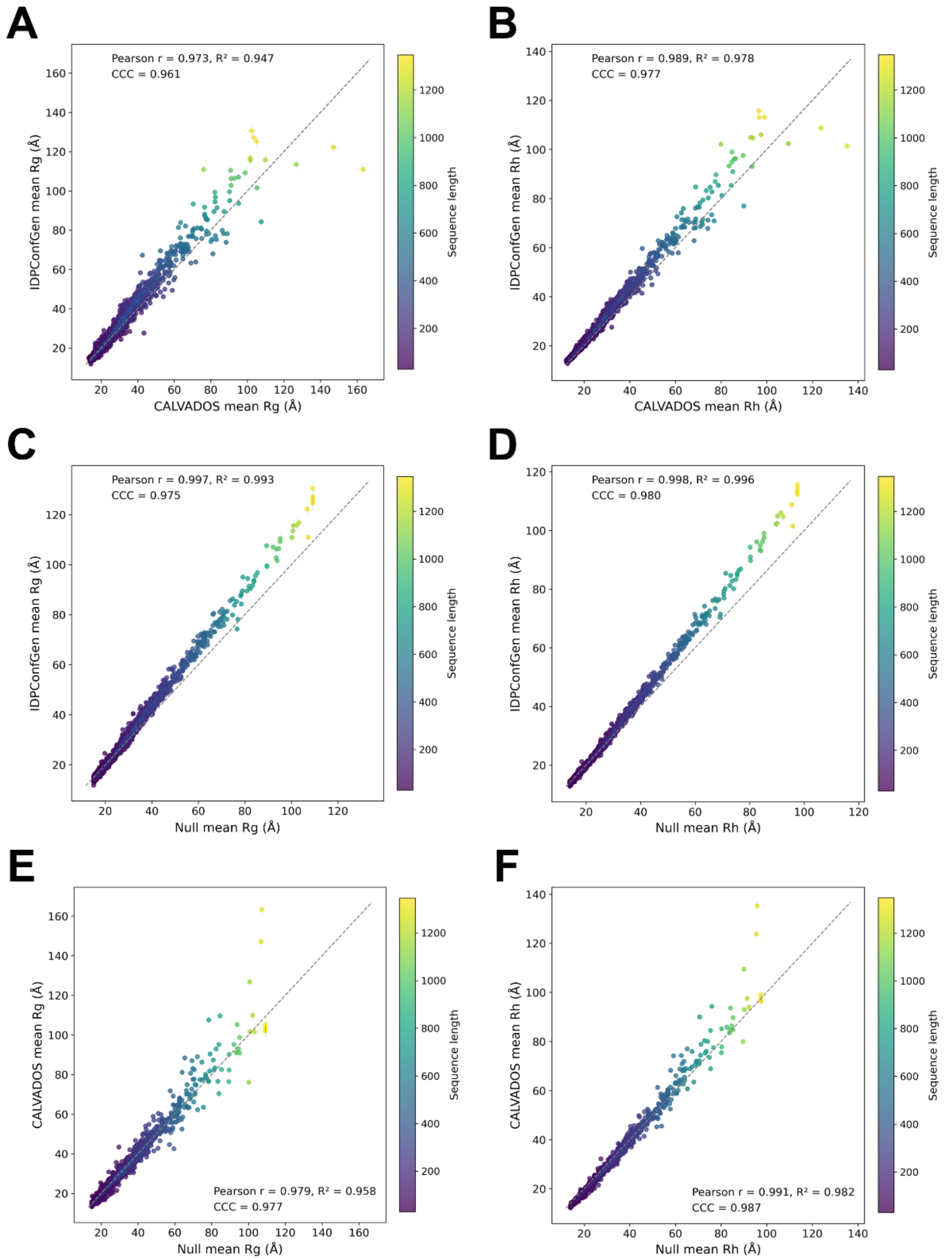


**Supplemental Figure S5. Scatter plots of mean $R_g$ and $R_h$ comparisons for 1,387 matching IDR sequences between CALVADOS and IDPConformerGenerator ensembles when generated in isolation.** (**A**) Mean $R_g$ of N = 100 conformers. Pearson r = 0.973, $R^2$ = 0.947, concordance correlation coefficient (CCC) = 0.961. (**B**) Mean $R_h$ of N = 100 conformers). Pearson r = 0.989, $R^2$ = 0.978, CCC = 0.977. (**C**) IDPCG mean $R_g$ of N = 100 conformers compared to a theoretical $R_g$ scaling law[4] of $R_g = 2.54 \times N^{0.522}$. (**D**) IDPCG mean $R_h$ of N = 100 conformers compared to a theoretical $R_h$ scaling law[5] of $R_h = 2.49 \times N^{0.509}$. (**E**) CALVADOS mean $R_g$ of N = 100 conformers compared to a theoretical $R_g$ scaling law[4] of $R_g = 2.54 \times N^{0.522}$. (**F**) CALVADOS mean $R_h$ of N = 100 conformers compared to a theoretical $R_h$ scaling law[5] of $R_h = 2.49 \times N^{0.509}$.

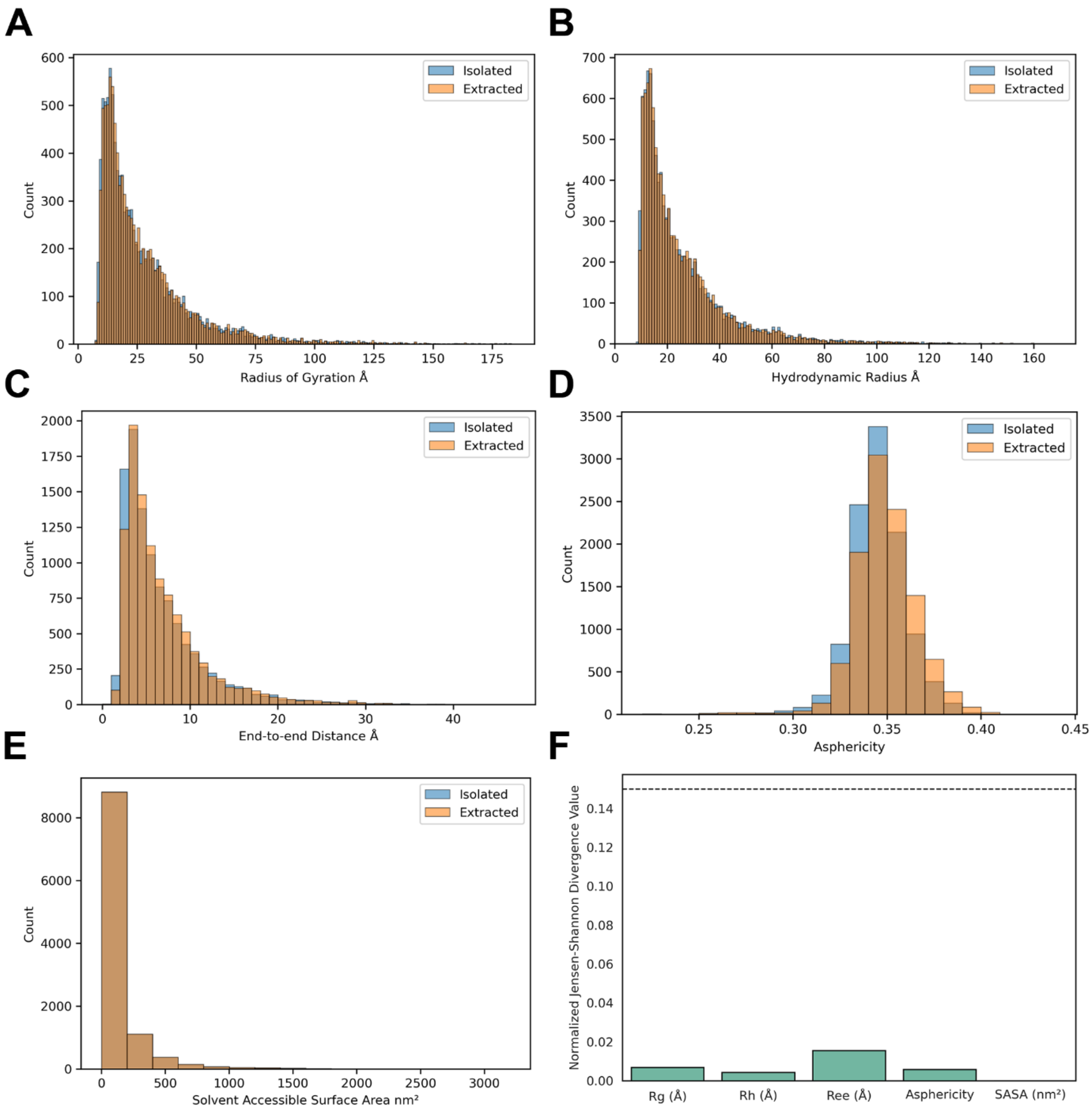


**Supplemental Figure S6.1. Distributions of ensemble structural properties (category 1, N = 10,698) between IDRs generated by IDPConformerGenerator in isolation and IDRs extracted from AFX-IDPCG ensembles.** Blue distributions represent IDR ensembles generated in isolation as IDPs by IDPConformerGenerator. Orange distributions represent IDRs extracted from the context of the folded domain from the AlphaFlex workflow. (**A**) $R_g$ (Å) with 1 Å bin sizes. (**B**) $R_h$ (Å) with 1 Å bin sizes. (**C**) $R_{ee}$ (Å) with 1 Å bin sizes. (**D**) Asphericity with bin sizes of 0.01 units. (**E**) SASA with 20 $nm^2$ bin sizes. (**F**) Normalized Jensen-Shannon (JS) statistics of all structural properties. Black dashed line represents a 0.15 cutoff for significantly different properties.

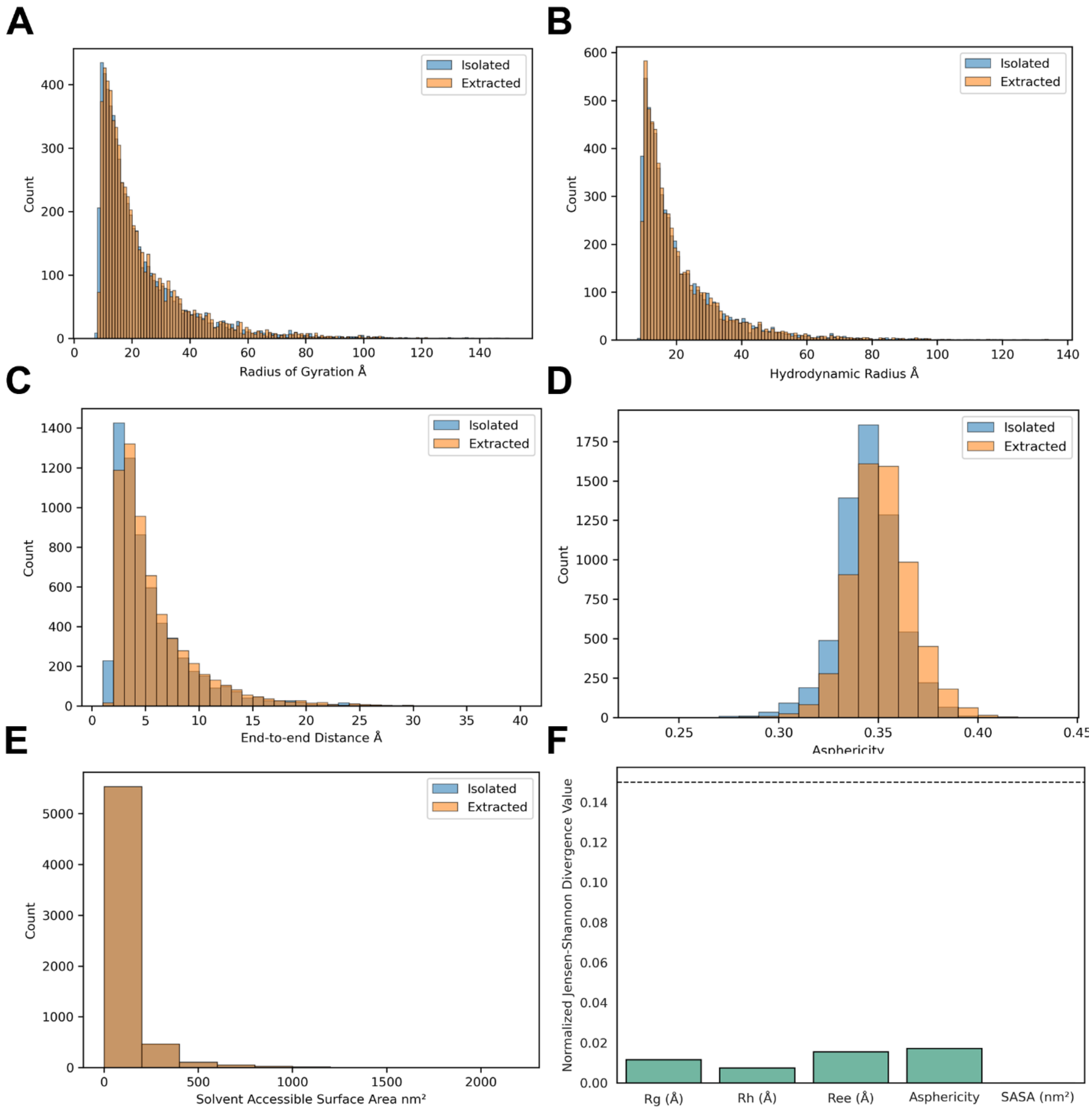


**Supplemental Figure S6.2. Distributions of ensemble structural properties (category 2, N = 6,226) between IDRs generated by IDPConformerGenerator in isolation and IDRs extracted from AFX-IDPCG ensembles.** Blue distributions represent IDR ensembles generated in isolation as IDPs by IDPConformerGenerator. Orange distributions represent IDRs extracted from the context of the folded domain from the AlphaFlex workflow. (**A**) $R_g$ (Å) with 1 Å bin sizes. (**B**) $R_h$ (Å) with 1 Å bin sizes. (**C**) $R_{ee}$ (Å) with 1 Å bin sizes. (**D**) Asphericity with bin sizes of 0.01 units. (**E**) SASA with 20 $nm^2$ bin sizes. (**F**) Normalized Jensen-Shannon (JS) statistics of all structural properties. Black dashed line represents a 0.15 cutoff for significantly different properties.

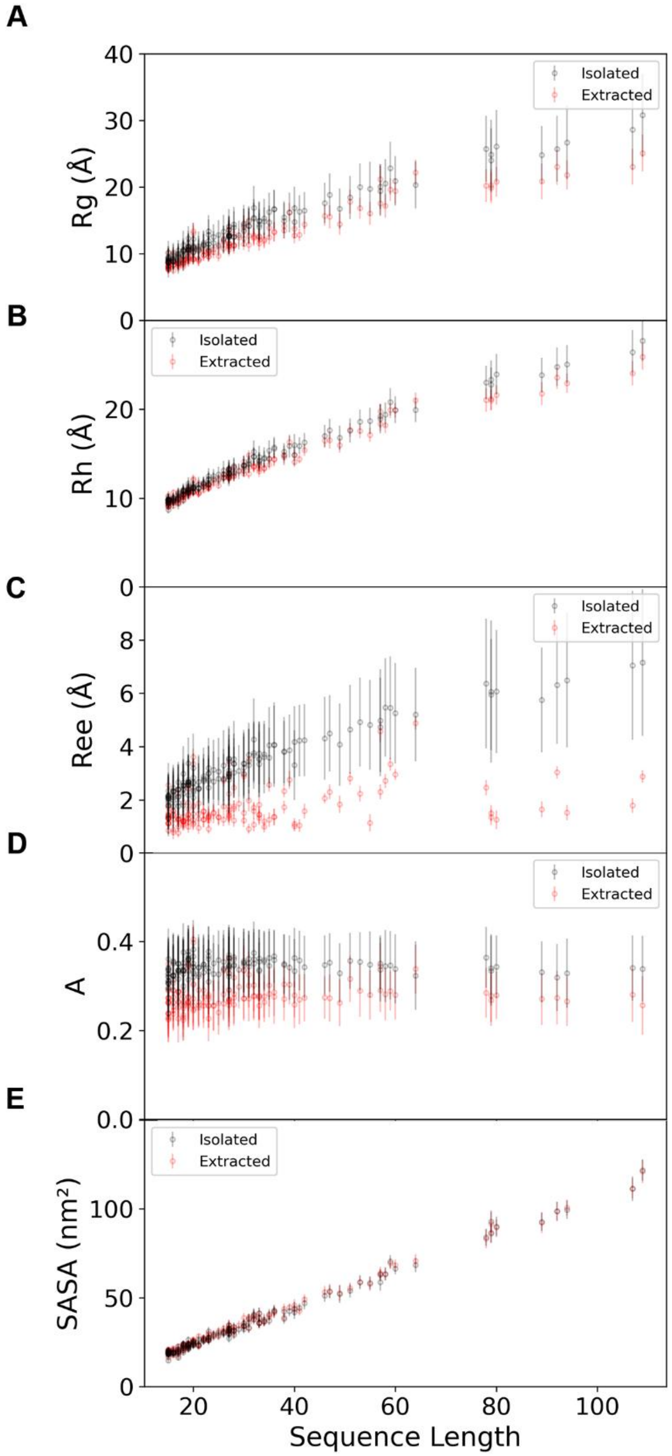


**Supplemental Figure S7. Scatter plot of different global structural properties against sequence length for category 3 AFX-IDPCG structures with a single loop IDR (N = 147 proteins).** IDRs generated by IDPConformerGenerator in isolation in black. IDRs extracted from AFX-IDPCG ensembles are in red. Lines for each dot represent the mean standard error for the ensembles (N = 100 conformers per protein. Units for A-C are in Angstroms (Å) and E in squared-nanometers ($nm^2$). (**A**) Radius of gyration ($R_g$). (**B**) Hydrodynamic radius ($R_h$). (**C**) End-to-end distance ($R_{ee}$). (**D**) Asphericity (A). (**E**) Solvent accessible surface area (SASA).

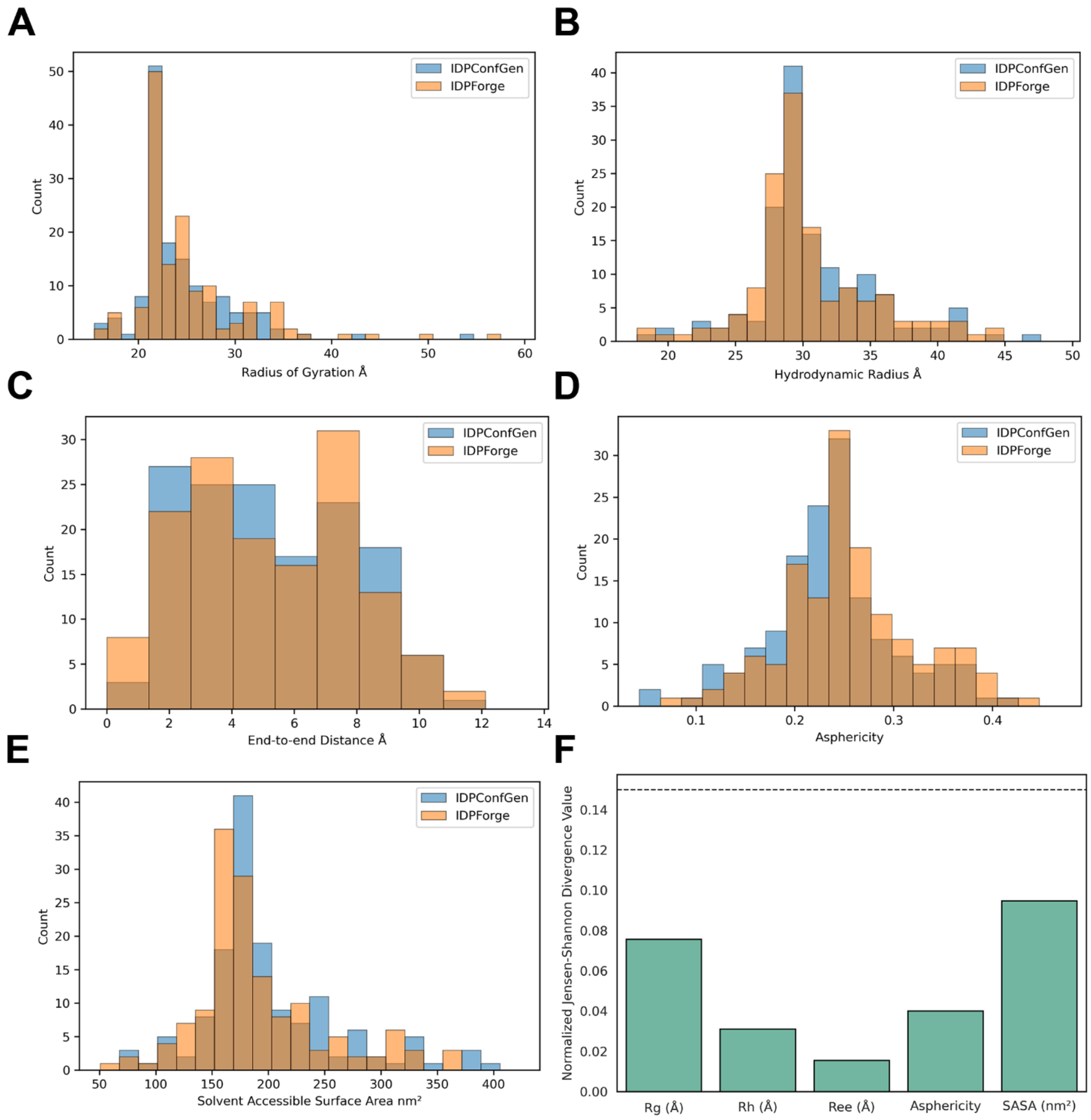


**Supplemental Figure S8. Distributions of ensemble structural properties (category 3, single loop IDR, N = 147 proteins) between full-length AFX-IDPCG and AFX-IDPForge ensembles.** Blue distributions represent full-length AlphaFlex-IDPConformerGenerator ensembles. Orange distributions represent full-length AlphaFlex-IDPForge ensembles. A) $R_g$ (Å) with 1.4 Å bin sizes. B) $R_h$ (Å) with 1.4 Å bin sizes. C) $R_{ee}$ (Å) with 1.3 Å bin sizes. D) Asphericity with bin sizes of 0.02 units. E) SASA with 1.6 $nm^2$ bin sizes. F) Normalized Jensen-Shannon (JS) statistics of all structural properties. Black dashed line represents a 0.15 cutoff for significantly different properties.

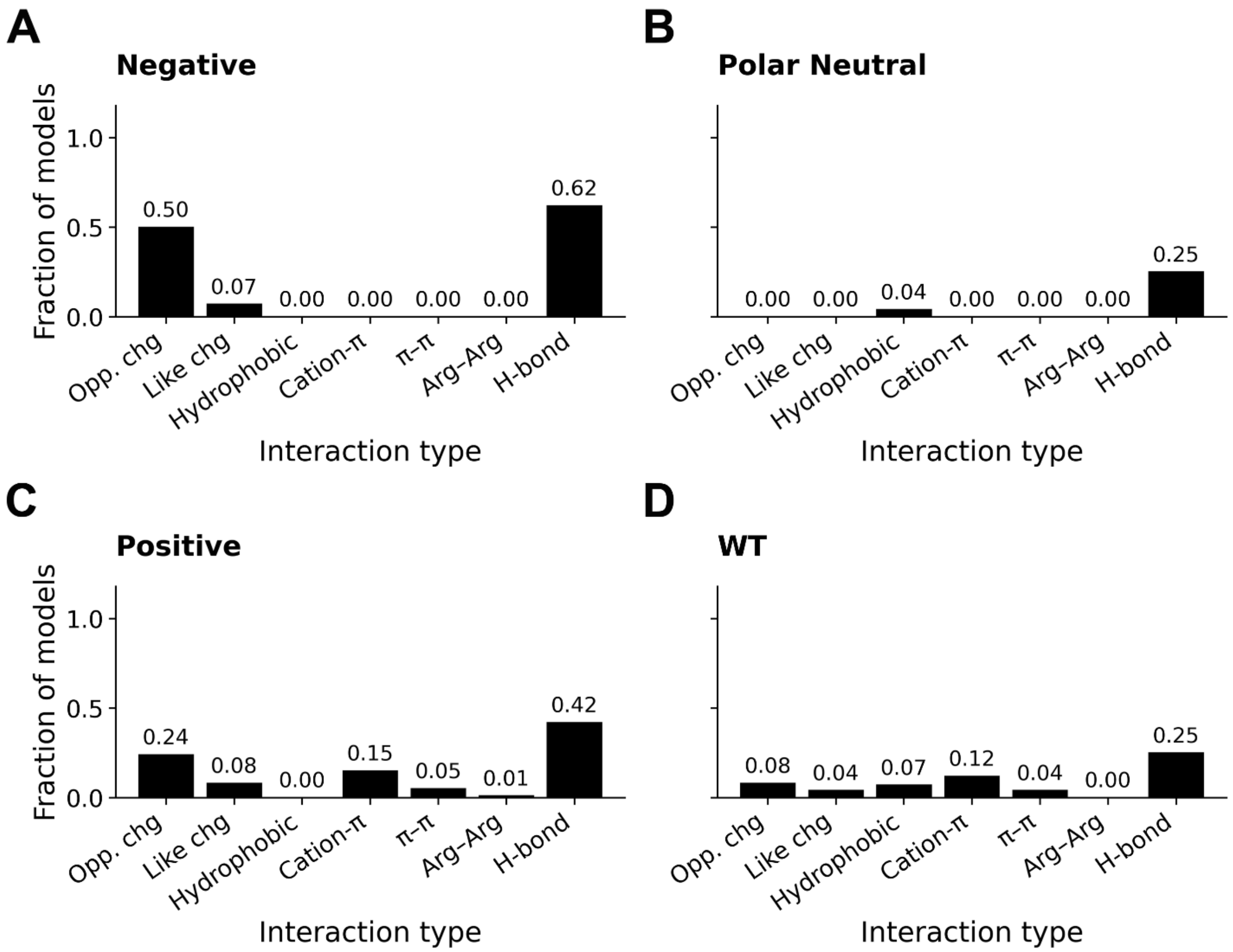


**Supplemental Figure S9.1. Sampling statistics for IDR:folded-domain interactions from chimeras of AFX-IDPCG (N = 100 conformers) ensembles for A6NI47.** X-Axis describes the type of interaction each ensemble contains. Fraction of models within the ensemble showing each contact type. Opp. chg is opposite-charge interactions. Like chg is like-charge interactions. (**A**) Net-negative N-terminal IDR. (**B**) Polar-neutral N-terminal IDR. (**C**) Net-positive N-terminal IDR. (**D**) Wildtype N-terminal IDR.

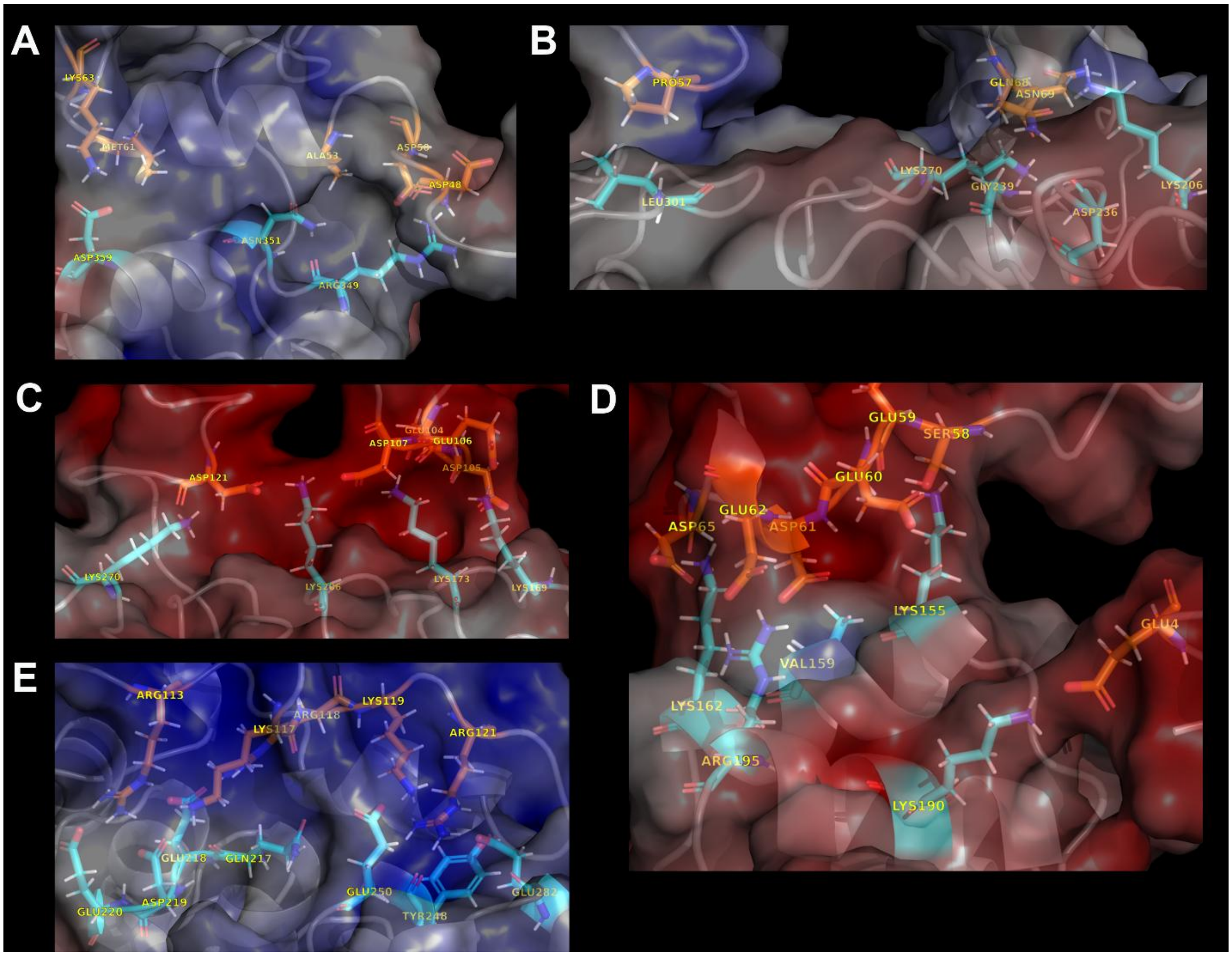


**Supplemental Figure S9.2. Contacts between the N-terminal IDR (residues 1-132) and the folded domain for A6NI47 in the chimera experiment.** Transparent electrostatic surfaces of the folded domain were generated with the PyMOL APBS electrostatics plugin[6]. Red surfaces are negative, blue surfaces are positive, grey surfaces are neutral. The interacting IDR (orange) and folded domain (cyan) residues. Refer to Supplemental Table S4 for the residue ranges of positive and negative folded domain patches. (**A**) WT N-terminal IDR. Two opposite-charge pairs: D50-R349, K63-D359, and six hydrogen-bond type pairs: D480-R349, D50-R349, A53-N351, M61-Q355, M61-D359, K63-D359. (**B**) Polar-neutral N-terminal IDR. One hydrophobic pair P57-L301 and four hydrogen-bond-like pairs Q68-K206; Q68-D236; Q68-G239; N69-K270 are observed. (**C**) and (**D**) are the net-negatively charged N-terminal IDR at different contact locations making electrostatic contacts on and near positive patches #1 and #2. Nine opposite-charge pairs: E60-K155; D61-K162; E62-K162; D65-K162; D105-K169; D105-K173; D107-K173; D107-K206; D121-K206, and 14 hydrogen-bond-pairs: E4-K190; S58-K155; E59-K155; D61-V159; D61-K162; E62-R195; D65-K162; E104-K173; D105-K169; E106-K169; D107-K173; D107-K206; D121-K206; D121-K270. (**E**) Net-positively charged N-terminal IDR. Five opposite-charge pairs: R113-E218; R113-D219; K117-E218; R118-E218; R121-E282, one cation-π: R121-Y248, one π-π: R121-Y248, and 13 hydrogen-bond-like pairs: R113-E218; R113-D219; R113-E220; K116-E250; K117-Q217; K117-E218; K117-D219; R118-C216; R118-E218; K119-E250; R121-Y248; R121-N249; R121-E282 are observed.

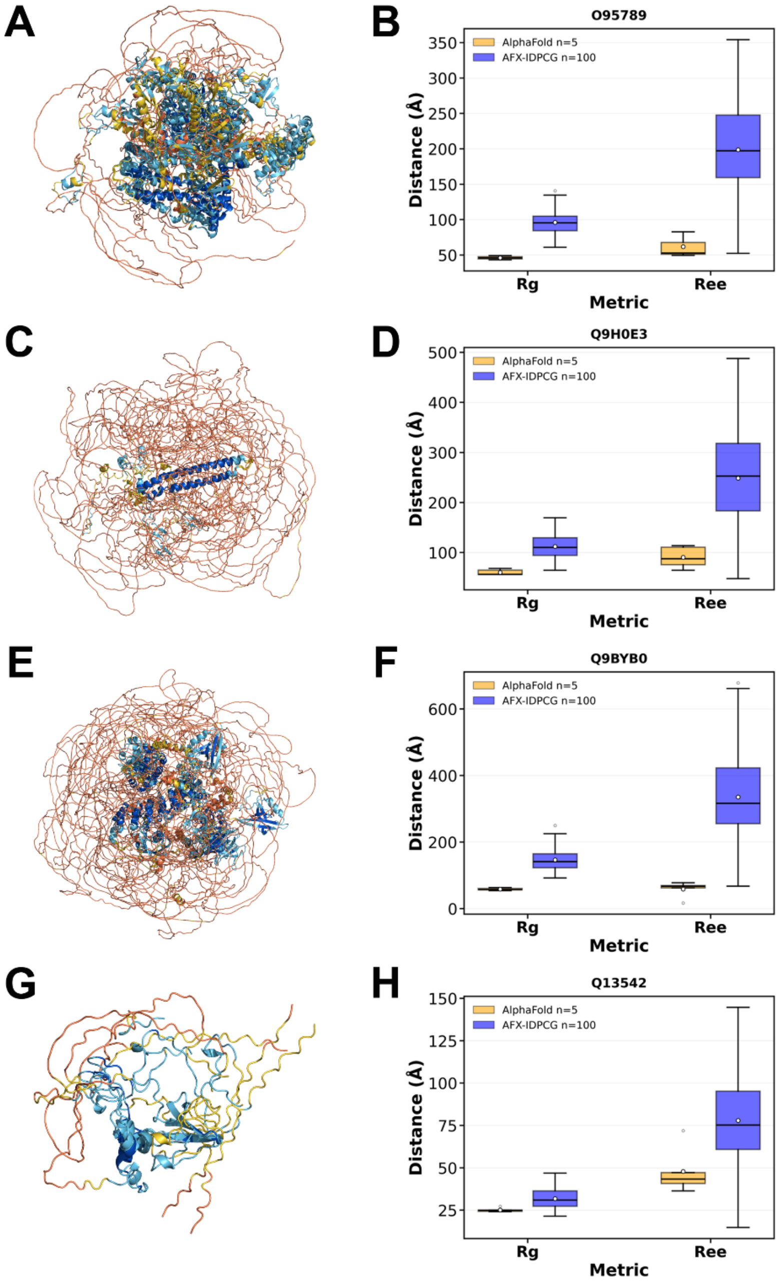


**Supplemental Figure S10.1. AlphaFold2 conformations (N = 5) of category 1, 2, and 3 proteins.** Conformations are colored by AlphaFold pLDDT. All 5 conformations are overlaid on top of each other aligned by their largest confidently predicted region. Predictions were generated by AlphaFold2 ColabFold[7]. (**A**) AlphaFold2 predictions of ZMYM6 (UniProt ID O95789), category 2 (**B**) Distribution of $R_g$ and $R_{ee}$ of AlphaFold2 (orange) and AFX-IDPCG (blue) ensembles for ZMYM6. (**C**) AlphaFold2 predictions of SAP130 (UniProt ID Q9H0E3), category 2 (**D**) Distribution of $R_g$ and $R_{ee}$ of AlphaFold2 (orange) and AFX-IDPCG (blue) ensembles for SAP130. (**E**) AlphaFold2 predictions of SHANK3 (UniProt ID Q9BYB0), category 3 (**F**) Distribution of $R_g$ and $R_{ee}$ of AlphaFold2 (orange) and AFX-IDPCG (blue) ensembles for SHANK3. (**G**) AlphaFold2 predictions of 4E-BP2 (UniProt ID Q13542), category 1 (**H**) Distribution of $R_g$ and $R_{ee}$ of AlphaFold2 (orange) and AFX-IDPCG (blue) ensembles for 4E-BP2.

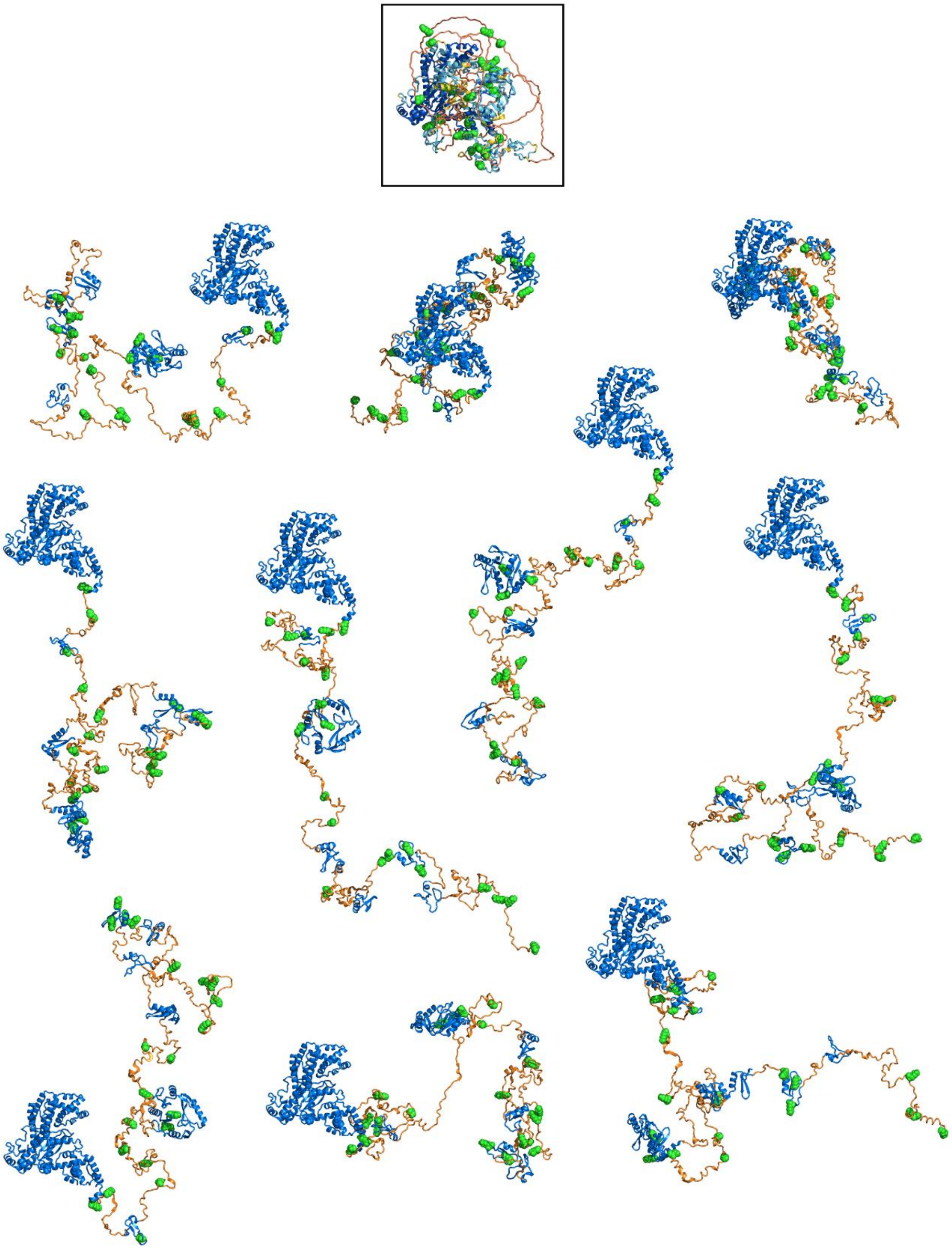

**Supplemental Figure S10.2. AlphaFlex conformations (N = 10) of ZMYM6 (UniProt ID O95789) highlighting residues that undergo post-translational modifications.** Residues that undergo PTMs defined by UniProt's PTM/Processing are modeled as green spheres. Topmost structure is the AFDB model of ZMYM6 outlined in a black box, colored by AlphaFold's definition of pLDDT with yellow and orange representing low (50 < pLDDT < 70) and very low (pLDDT < 50) confidence, respectively, and light blue (70 < pLDDT < 90) and blue (90 < pLDDT) representing high and very confidence, respectively[2]. Bottom 10 AlphaFlex conformations are the same scale and orientation. IDRs are colored orange while folded domains are blue for the AlphaFlex conformers.

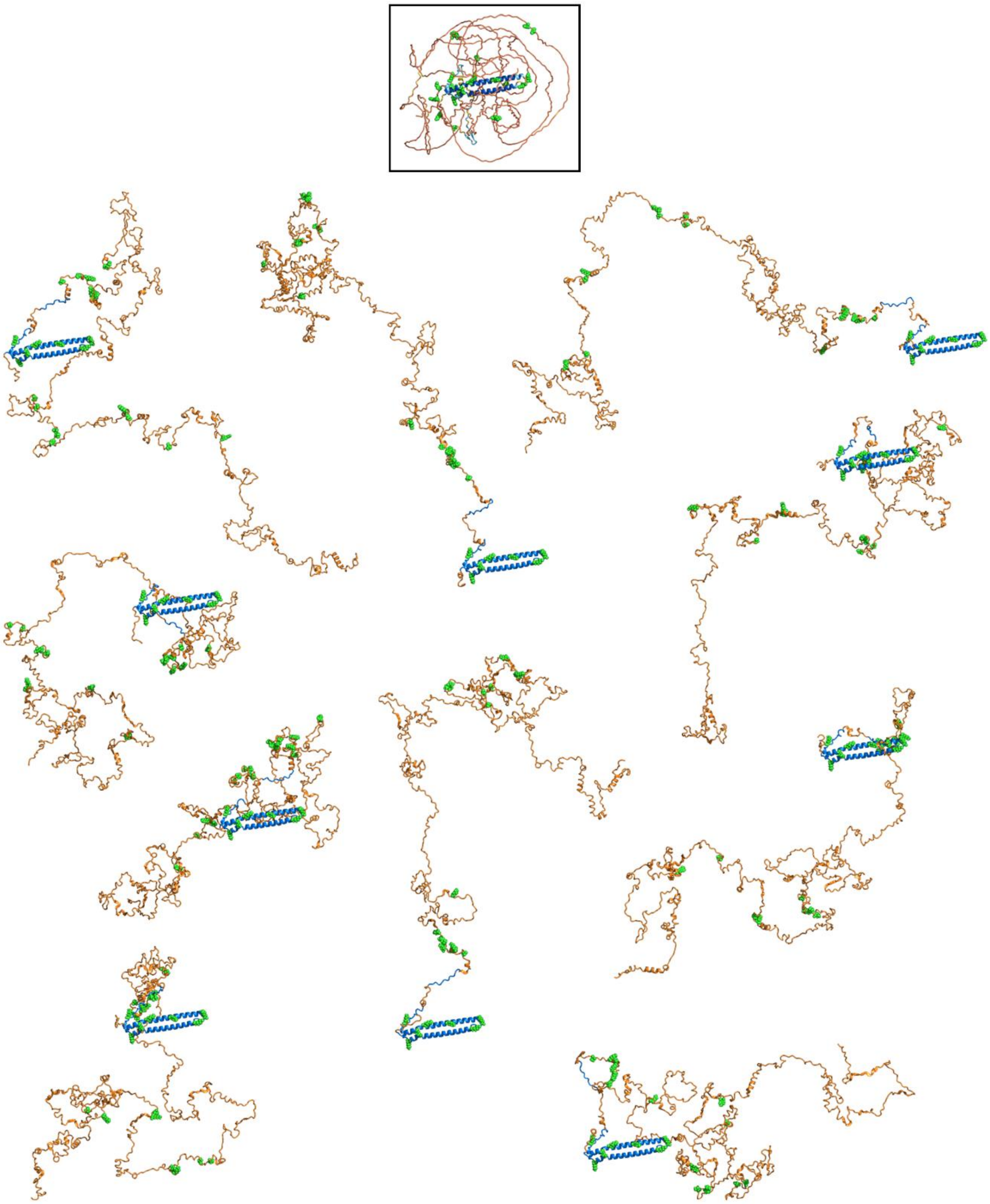

**Supplemental Figure S10.3. AlphaFlex conformations (N = 10) of SAP130 (UniProt ID Q9H0E3) highlighting residues that undergo post-translational modifications.** Residues that undergo PTMs defined by UniProt's PTM/Processing are modeled as green spheres. Topmost structure is the AFDB model of SAP130 outlined in a black box, colored by AlphaFold's definition of pLDDT with yellow and orange representing low (50 < pLDDT < 70) and very low (pLDDT < 50) confidence, respectively, and light blue (70 < pLDDT < 90) and blue (90 < pLDDT) representing high and very confidence, respectively[2]. Bottom 10 AlphaFlex conformations are the same scale and orientation. IDRs are colored orange while folded domains are blue for the AlphaFlex conformers.

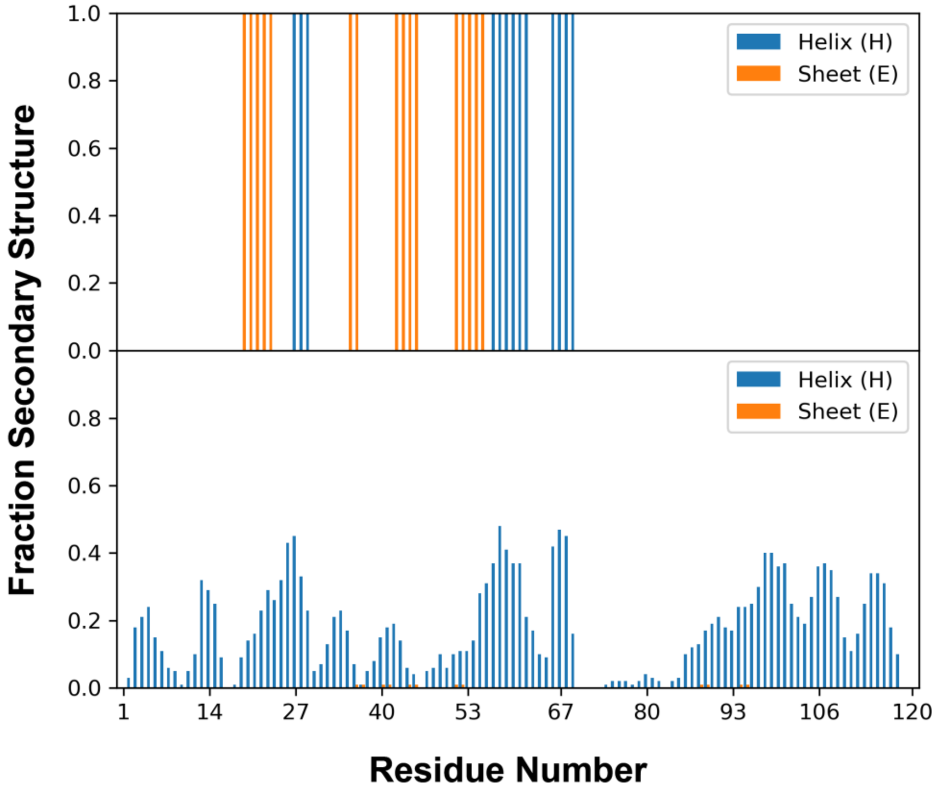


**Supplemental Figure S11. Fractional secondary structure from AlphaFold (top) compared to AlphaFlex ensemble (bottom) of 4E-BP2 (UniProt ID Q13542).** Fractional DSSP L is hidden for visualization purposes. N = 100 AFX-IDPCG ensembles of 4E-BP2 analyzed with DSSP for the bottom panel. Fractional α-helical structure is present in the canonical binding α-helix of 4E-BP2 from residue 54-65, with β-strands. The formation of the β-structure under phosphorylated conditions sequesters the canonical binding α-helix, however the AFDB structure contains a combination of α-helices and β-strands in the residue ranges for the binding α-helix, confusing the mechanistic insights.

**Supplemental Table S1.1. Distribution divergence metrics for sequence-matched CALVADOS and IDPConformerGenerator distributions (isolated IDRs).** *p* represents the distribution of CALVADOS ensembles[8]; *q* represents the distribution of IDPConformerGenerator ensembles. N = 1,387. Statistical measurements of KL and JS are unitless.

| Global Structural Property | $KL_{(\tilde{p}\|\|\tilde{q})}$ | $KL_{(\tilde{q}\|\|\tilde{p})}$ | $JS_{(\tilde{p},\tilde{q})}$ | $JS_{norm}(\tilde{p},\tilde{q})$ |
|---|---|---|---|---|
| Radius of gyration | 0.067 | 0.130 | 0.009 | 0.012 |
| Hydrodynamic radius | 0.068 | 0.190 | 0.009 | 0.013 |
| End-to-end distance | 0.088 | 0.253 | 0.014 | 0.021 |
| Asphericity | 2.480 | 0.639 | 0.149 | 0.215 |
| Solvent accessible surface area | 0.076 | 0.021 | 0.005 | 0.088 |

**Supplemental Table S1.2. Distribution divergence metrics for extracted and isolated ensembles from AlphaFlex category 1.** *p* represents the distribution of IDR ensembles generated in isolation by IDPConformerGenerator for category 1; *q* represents the distribution of AFX-IDPCG IDR ensembles extracted from the context of the folded domain for category 1. N = 10,698. Statistical measurements of KL and JS are unitless.

| Global Structural Property | $KL_{(\tilde{p}\|\|\tilde{q})}$ | $KL_{(\tilde{q}\|\|\tilde{p})}$ | $JS_{(\tilde{p},\tilde{q})}$ | $JS_{norm}(\tilde{p},\tilde{q})$ |
|---|---|---|---|---|
| Radius of gyration | 0.064 | 0.057 | 0.005 | 0.007 |
| Hydrodynamic radius | 0.038 | 0.053 | 0.003 | 0.004 |
| End-to-end distance | 0.018 | 0.045 | 0.004 | 0.006 |
| Asphericity | 0.044 | 0.083 | 0.012 | 0.017 |
| Solvent accessible surface area | 0.000 | 0.000 | 0.000 | 0.000 |

**Supplemental Table S1.3. Distribution divergence metrics for extracted and isolated ensembles from AlphaFlex category 2.** *p* represents the distribution of IDR ensembles generated in isolation by IDPConformerGenerator for category 2; *q* represents the distribution of AFX-IDPCG IDR ensembles extracted from the context of the folded domain for category 2. N = 6,226. Statistical measurements of KL and JS are unitless.

| Global Structural Property | $KL_{(\tilde{p}\|\|\tilde{q})}$ | $KL_{(\tilde{q}\|\|\tilde{p})}$ | $JS_{(\tilde{p},\tilde{q})}$ | $JS_{norm}(\tilde{p},\tilde{q})$ |
|---|---|---|---|---|
| Radius of gyration | 0.118 | 0.083 | 0.008 | 0.011 |
| Hydrodynamic radius | 0.086 | 0.078 | 0.005 | 0.007 |
| End-to-end distance | 0.078 | 0.069 | 0.012 | 0.017 |
| Asphericity | 0.153 | 0.112 | 0.023 | 0.037 |
| Solvent accessible surface area | 0.000 | 0.000 | 0.000 | 0.000 |

**Supplemental Table S1.4. Distribution divergence metrics for extracted and isolated ensembles from AlphaFlex category 3 proteins with a single loop IDR.** *p* represents the distribution of IDR ensembles generated in isolation by IDPConformerGenerator for category 3; *q* represents the distribution of AFX-IDPCG IDR ensembles extracted from the context of the folded domain for category 3. N = 147. Statistical measurements of KL and JS are unitless.

| Global Structural Property | $KL_{(\tilde{p}\|\|\tilde{q})}$ | $KL_{(\tilde{q}\|\|\tilde{p})}$ | $JS_{(\tilde{p},\tilde{q})}$ | $JS_{norm}(\tilde{p},\tilde{q})$ |
|---|---|---|---|---|
| Radius of gyration | 1.338 | 0.704 | 0.070 | 0.010 |
| Hydrodynamic radius | 0.805 | 0.450 | 0.025 | 0.036 |
| End-to-end distance | 5.492 | 5.492 | 0.330 | 0.473 |
| Asphericity | 3.668 | 20.985 | 0.455 | 0.657 |
| Solvent accessible surface area | 0.211 | 0.211 | 0.010 | 0.014 |

**Supplemental Table S1.5. Distribution divergence metrics for full-length AlphaFlex category 3 ensembles generated with IDPConformerGenerator and IDPForge.** *p* represents the distribution of AFX-IDPCG ensembles for category 3; q represents the distribution of AFX-IDPForge ensembles for category 3 with a single loop IDR. N = 147 proteins. Statistical measurements of KL and JS are unitless.

| Global Structural Property | $KL_{(\tilde{p}\|\|\tilde{q})}$ | $KL_{(\tilde{q}\|\|\tilde{p})}$ | $JS_{(\tilde{p},\tilde{q})}$ | $JS_{norm}(\tilde{p},\tilde{q})$ |
|---|---|---|---|---|
| Radius of gyration | 1.647 | 1.245 | 0.052 | 0.076 |
| Hydrodynamic radius | 0.259 | 0.444 | 0.021 | 0.031 |
| End-to-end distance | 0.041 | 0.046 | 0.011 | 0.015 |
| Asphericity | 0.466 | 0.455 | 0.028 | 0.040 |
| Solvent accessible surface area | 1.146 | 0.986 | 0.067 | 0.095 |

**Supplemental Table S1.6. Distribution divergence metrics for IDRs extracted from AlphaFlex category 3 ensembles generated with IDPConformerGenerator and IDPForge.** *p* represents the distribution of IDRs extracted AFX-IDPCG ensembles for category 3; *q* represents the distribution of IDRs extracted AFX-IDPForge ensembles for category 3. N = 147. Note that these calculations took IDPConformerGenerator 3 weeks normalized to a single machine with 50 CPU workers while it took IDPForge less than 1 week normalized to a single machine with 1 Nvidia A40 GPU. Statistical measurements of KL and JS are unitless.

| Global Structural Property | $KL_{(\tilde{p}\|\|\tilde{q})}$ | $KL_{(\tilde{q}\|\|\tilde{p})}$ | $JS_{(\tilde{p},\tilde{q})}$ | $JS_{norm}(\tilde{p},\tilde{q})$ |
|---|---|---|---|---|
| Radius of gyration | 2.392 | 0.553 | 0.096 | 0.138 |
| Hydrodynamic radius | 1.693 | 0.207 | 0.052 | 0.075 |
| End-to-end distance | 0.784 | 0.789 | 0.063 | 0.091 |
| Asphericity | 1.827 | 15.564 | 0.320 | 0.461 |
| Solvent accessible surface area | 1.482 | 0.770 | 0.059 | 0.085 |

**Table 2.1 Benchmarks on quality-controlled SAXS curves from the SASBDB for full-length proteins.** Lower $\chi^2/N$ represents better fit with experimental data. Pepsi-SAXS v3.0[9] used to calculate the predicted SAXS curves from structures. The mean predicted I(q) of N = 100 conformers for AF-CALVADOS and AFX-IDPCG were used to fit the experimental curve.

| UniProt ID | Category | Protein Length | % IDR | AFDB ($\chi^2/N$) | AF-CALVADOS ($\chi^2/N$) | AFX-IDPCG ($\chi^2/N$) |
|---|---|---|---|---|---|---|
| P26599 | 3 | 531 | 36% | 7.237 | 0.514 | 0.593 |
| P54252 | 3 | 361 | 55% | 31.696 | 0.475 | 0.847 |
| P54727 | 2 | 409 | 53% | 19.249 | 0.759 | 1.578 |
| P98170 | 2 | 497 | 32% | 90.480 | 0.857 | 1.211 |
| Q99418 | 2 | 400 | 15% | 130.251 | 1.678 | 1.580 |
| Q9NRC8 | 3 | 400 | 32% | 2.077 | 1.354 | 1.046 |
| Q9NRR5 | 2 | 601 | 78% | 53.716 | 0.977 | 1.515 |
| Q9Y3A5 | 2 | 250 | 26% | 4.189 | 0.593 | 1.449 |
| | | | **Average $\chi^2/N$** | 42.362 | 0.901 | 1.228 |

**Table 2.2 Benchmarks on NMR chemical shift data from the BMRB for the extracted IDRs in the context of the full-length protein.** Lower $\chi^2/N$ represents better fit with experimental data. UCBShift v2.0[10] used to calculate the predicted chemical shifts from structures. The mean predicted chemical shifts of each atom of N = 100 conformers for AF-CALVADOS and AFX-IDPCG were used to fit the experimental chemical shifts. *Signifies that ensembles were not found in the AF-CALVADOS repository. AF-CALVADOS IDR-boundaries were not publicly available, so AlphaFlex IDR boundaries were used for AF-CALVADOS ensembles.

| UniProt ID | Category | Protein Length | % IDR | AFDB ($\chi^2/N$) | AF-CALVADOS ($\chi^2/N$) | AFX-IDPCG ($\chi^2/N$) |
|---|---|---|---|---|---|---|
| O76070 | 1 | 127 | 100% | 6.434 | 2.884 | 2.226 |
| P06730 | 1 | 217 | 24% | 2.567 | 2.224 | 3.822 |
| P37840 | 1 | 140 | 100% | 4.354 | 0.813 | 1.172 |
| P42771 | 1 | 156 | 15% | 0.696 | 0.247 | 0.215 |
| P46109 | 2 | 303 | 26% | 7.995 | 7.323 | 7.580 |
| Q00688 | 1 | 224 | 17% | 1.436 | 1.510 | 1.616 |
| Q13542 | 1 | 120 | 100% | 3.414 | 3.167 | 2.907 |
| Q15004 | 1 | 111 | 100% | 1.738 | 1.080 | 1.200 |
| Q15121 | 1 | 130 | 66% | 2.189 | 1.998 | 2.747 |
| Q15185 | 1 | 160 | 34% | 2.229 | 0.861 | 1.086 |
| Q96I45 | 1 | 108 | 53% | 3.139 | * | 2.643 |
| Q9P298 | 2 | 99 | 52% | 9.832 | * | 8.187 |
| Q9Y237 | 1 | 131 | 31% | 7.372 | 6.192 | 5.050 |
| | | | **Average $\chi^2/N$** | 4.107 | 2.573 | 3.112 |

**Supplemental Table S3.1. Percentage of DSSP secondary structure.** IDPConformerGenerator database which includes RCSB PDB X-Ray crystal structures with resolution better than or equal to 2.0 Å as of 2024 are given as a reference. AlphaFold pLDDT < 70 across different thresholds of minimum residue length (N = 14,792 proteins). CALVADOS representations of IDRs after reducing the number of conformations to 100 per protein (N = 28, 058 IDRs). IDRs extracted from the AFX-IDPForge ensembles (N = 203). IDRs extracted from the AFX-IDPCG ensembles (N = 17,071 IDRs). Values with an * do not have an associated mean standard deviation since they are not ensembles. **Only the experimentally benchmarked AF-CALVADOS (N = 19) ensembles were calculated here. AF-CALVADOS IDR-boundaries were not publicly available, so AlphaFlex IDR boundaries were used for AF-CALVADOS ensembles, likely explaining the presence of helical and beta structure.

| *Method* | *% H (α-helical) ± STD* | *% E (extended β) ± STD* | *% L (loop) ± STD* |
|---|---|---|---|
| *RCSB PDB Database* | 35.4* | 23.0* | 41.6* |
| *AFDB* | 10.3* | 0.8* | 88.9* |
| *CALVADOS* | 0.1 ± 0.7 | 0.0 ± 0.0 | 99.9 ± 0.7 |
| *AF-CALVADOS*** | 7.7 ± 12.4 | 1.4 ± 3.5 | 90.9 ± 12.6 |
| *AFX-IDPForge* | 2.9 ± 6.4 | 0.2 ± 1.7 | 96.9 ± 6.6 |
| *AFX-IDPCG* | 23.4 ± 14.1 | 0.1 ± 0.8 | 75.5 ± 14.1 |

**Supplemental Table S3.2. Mean percentage ± standard deviation of DSSP secondary structure of extracted IDRs from the AFX-IDPForge ensembles across different thresholds of minimum residue length (N = 203 ensembles).**

| *Minimum Length* | *% H (α-helical) ± STD* | *% E (extended β) ± STD* | *% L (loop) ± STD* |
|---|---|---|---|
| *1* | 2.9 ± 6.4 | 0.2 ± 1.7 | 96.9 ± 6.6 |
| *50* | 3.3 ± 4.4 | 0.2 ± 1.3 | 96.5 ± 4.5 |
| *100* | 3.2 ± 3.8 | 0.2 ± 1.8 | 96.6 ± 3.8 |
| *250* | 2.6 ± 2.0 | 0.3 ± 0.6 | 97.1 ± 2.1 |

**Supplemental Table S3.3. Mean percentage ± standard deviation of DSSP secondary structure of extracted IDRs from the AFX-IDPCG ensembles across different thresholds of minimum residue length (N = 17,071 ensembles).**

| *Minimum Length* | *% H (α-helical) ± STD* | *% E (extended β) ± STD* | *% L (loop) ± STD* |
|---|---|---|---|
| *1* | 23.4 ± 14.1 | 0.1 ± 0.8 | 75.5 ± 14.1 |
| *50* | 24.1 ± 10.0 | 0.1 ± 0.5 | 75.9 ± 10.0 |
| *100* | 24.2 ± 8.5 | 0.1 ± 0.4 | 75.7 ± 8.5 |
| *500* | 24.8 ± 6.7 | 0.1 ± 0.2 | 75.1 ± 6.7 |
| *1000* | 22.9 ± 5.7 | 0.1 ± 0.2 | 77.0 ± 5.7 |

**Supplemental Table S3.4. Net percentage of DSSP secondary structure of AlphaFold pLDDT < 70 across different thresholds of minimum residue length (N = 14,792 proteins).**

| *Minimum Length* | *% H (α-helical) ± STD* | *% E (extended β) ± STD* | *% L (loop) ± STD* |
|---|---|---|---|
| *1* | 10.3 | 0.8 | 88.9 |
| *50* | 7.5 | 0.7 | 91.8 |
| *100* | 5.9 | 0.7 | 93.4 |
| *500* | 2.1 | 0.5 | 97.3 |
| *1000* | 1.3 | 0.8 | 97.93 |

**Supplemental Table S4. Charged folded domain patches as identified by the AlphaFlex surface chemistry profiler for A6NI47.**

| *Patch Number* | *Net Charge* | *Residue Numbers* | *Sequence* |
|---|---|---|---|
| *1* | +7 | 136, 137,142,145,151,154,155 | RRKRKRK |
| *2* | +6 | 168,169,171,173,205,206 | KKKKKK |
| *3* | +6 | 252,256,289,293,295 | KKKKKK |
| *4* | +3 | 190,195,196 | KRR |
| *5* | +3 | 268,270,303 | KKR |
| *6* | -3 | 250,251,282 | EDE |

**Supplemental Table S5. Hardware specifications of the computational resources used to calculate the AlphaFlex database.** DRAC represents the Digital Research Alliance of Canada clusters, where the CPU and RAM listed are only the resources requested from the cluster. Similarly, Savio is the compute cluster located at the University of Berkeley, California.

| *Sponsor* | *Machine* | *CPU* | *GPU* | *RAM* |
|---|---|---|---|---|
| *Julie Forman-Kay* | Niagara-DRAC<br>Cedar-DRAC<br>Nibi-DRAC | 50 threads | - | 64 GB |
| *Julie Forman-Kay* | Narwhal | AMD Threadripper 2990wx (32c/64t) | Nvidia RTX 3070 Ti (8 GB) | 256 GB |
| *Simon Sharpe* | Niagara-DRAC<br>Cedar-DRAC<br>Nibi-DRAC | 50 threads | - | 64 GB |
| *SickKids Molecular Medicine Program* | Haemonchus | 2x Intel Xeon Silver 4314 (16c/32t) | - | 320 GB |
| *Alan Moses* | Valrhona | 25 threads | - | 64 GB |
| *Hue Sun Chan* | node3-3<br>node3-4<br>node3-5 | 32 threads | 2x Nvidia RTX 2080 Ti (12 GB) | 64 GB |
| *Hue Sun Chan* | node2-9<br>node2-10<br>node2-11<br>node2-13<br>node2-14 | 16 threads | - | 32 GB |
| *Teresa Head-Gordon* | Savio | 16 threads | Nvidia A40 (48 GB) | 32 GB |
